# Phenotypic differences in reversible attachment behavior reveal distinct *P. aeruginosa* surface colonization strategies

**DOI:** 10.1101/798843

**Authors:** Calvin K. Lee, Jérémy Vachier, Jaime de Anda, Kun Zhao, Amy E. Baker, Rachel R. Bennett, Catherine R. Armbruster, Kimberley A. Lewis, Rebecca L. Tarnopol, Charles J. Lomba, Deborah A. Hogan, Matthew R. Parsek, George A. O’Toole, Ramin Golestanian, Gerard C. L. Wong

## Abstract

Despite possessing the machinery to sense, adhere to, and proliferate on surfaces, it is commonly observed that bacteria initially have a difficult time attaching to a surface. Before forming a bacterial biofilm, planktonic bacteria exhibit a random period of transient surface attachment known as “reversible attachment” which is poorly understood. Using community tracking methods at single-cell resolution, we examine how reversible attachment progresses during initial stages of surface sensing. *Pseudomonas aeruginosa* strains PAO1 and PA14, which exhibit similar exponential trends of surface cell population increase, show unanticipated differences when the behavior of each cell was considered at the full lineage level and interpreted using the unifying quantitative framework of an exactly solvable stochastic model. Reversible attachment comprises two regimes of behavior, processive and nonprocessive, corresponding to whether cells of the lineage stay on the surface long enough to divide, or not, before detaching. Stark differences between PAO1 and PA14 in the processive regime of reversible attachment suggest the existence of two complementary surface colonization strategies, which are roughly analogous to “immediate-” vs “deferred-gratification” in a prototypical cognitive-affective processing system. PAO1 lineages commit relatively quickly to a surface compared to PA14 lineages. PA14 lineages allow detaching cells to retain memory of the surface so that they are primed for improved subsequent surface attachment. In fact, it is possible to identify motility suppression events in PA14 lineages in the process of surface commitment. We hypothesize that these contrasting strategies are rooted in downstream differences between Wsp-based and Pil-Chp-based surface sensing systems.

## Importance

The initial pivotal phase of bacterial biofilm formation known as “reversible attachment,” where cells undergo a period of transient surface attachment, is at once universal and poorly understood. What is more, although we know that reversible attachment culminates ultimately in irreversible attachment, it is not clear how reversible attachment progresses phenotypically as bacterial surface sensing circuits fundamentally alters cellular behavior. We analyze diverse observed bacterial behavior one family at a time (defined as a full lineage of cells related to one another by division) using a unifying stochastic model and show that it leads to new insights on the time evolution of reversible attachment. Our results unify apparently disparate findings in the literature regarding early events in biofilm formation by PAO1 and PA14 strains.

## Introduction

Biofilms are surface-adhered communities or suspended aggregates of bacteria that have increased tolerance to environmental stresses and antibiotics, and impact human health and the environment in complex ways. These biofilms can be harmful by causing diseases (1, 2), and can be beneficial by serving as commensals in various hosts as well as having applications in bioremediation and energy production (3). A critical step in forming a bacterial biofilm is surface sensing (4), where free-swimming planktonic cells detect, attach to, and physiologically respond to a surface. Recent work has shown that different appendages or extracellular structures, such as flagella (5, 6) or type IV pili (TFP) (7, 8) are involved in activating cellular responses (e.g., protein production, motility, and biofilm formation) during surface sensing. In many bacterial species, these responses are primarily controlled by intracellular secondary messenger molecules, such as cyclic diguanylate (c-di-GMP) (9–16) and cyclic AMP (cAMP) (8, 17, 18). For the case of *Pseudomonas aeruginosa*, a clinically relevant model system (19), there are at least two well-studied but distinct surface sensing circuits, the Wsp and the Pil-Chp systems, that can contribute to initiating biofilm formation. In our current understanding, the Wsp system senses through the membrane-bound, chemosensory-like Wsp protein complex which localizes laterally along the cell body (10), activating the diguanylate cyclases WspR and c-di-GMP synthesis via a mechanism that requires clustering of (20). On the other hand, the Pil-Chp system senses a surface through polarly-localized TFP, which activates the adenylate cyclases CyaB and results in cAMP synthesis. Increased cAMP levels then induces the production and secretion of PilY1, which in turn activate the diguanylate cyclases SadC and results in c-di-GMP synthesis (17). Downstream consequences of c-di-GMP synthesis include exopolysaccharide (EPS) production and motility suppression. Different strains of *P. aeruginosa*, such as PAO1 and PA14, utilize these surface sensing mechanisms to varying extents. The PAO1 strain predominantly uses the Wsp system (21) leading to the surface deposition of the EPS Psl (22, 23), while PA14 predominantly uses the Pil-Chp system leading to the suppression of surface motility (17) and production of a Pel-dominant biofilm matrix (24).

Despite the existence of diverse machinery to sense, adhere to, and proliferate on surfaces, it is commonly observed that bacteria initially seem to have a difficult time attaching to a surface, as indicated by typical flow cell studies where *P. aeruginosa* often takes >20 h before attaching to the surface in large numbers (25, 26). This phenomenon was first reports in the 1930s (27, 28). Using high speed microscopy to measure the distribution of surface residence times, it was previously observed that the overwhelming majority of cells that land on the surface eventually detach, and it is only after a prolonged and variable time lag that cells begin to rapidly cover the surface (8). Reversible attachment is counterintuitive and difficult to understand for a number of reasons. We stress that the low apparent probability of successful attachment is not simply a matter of cells “bouncing” off the surface. (During reversible attachment, it is not uncommon for cells to attach and stay long enough to divide but then subsequently detach.) Moreover, the unpredictability of reversible attachment cannot be circumvented with better measurement statistics: the duration of reversible attachment always appears random and do not converge to a specific duration for the same initial conditions. This combination of characteristics in reversible attachment, low probability of success, intrinsic time dependence, and structurally random outcomes, suggests that use of a stochastic model may lead to new understanding. From a foundational perspective of surface sensing, although we know that reversible attachment can culminate in irreversible attachment, it is not clear how reversible attachment progresses phenotypically as bacterial surface sensing circuits fundamentally alters cellular behavior, and ultimately improve on an initial attachment probability of effectively zero.

Here, we show that the use of an exactly solvable “divide-detach” stochastic model, designed to examine the reversible attachment behaviors of *P. aeruginosa* PAO1 and PA14 lineages in the form of family trees, reveals differences in their biofilm formation behavior during reversible attachment. Within this model, reversible attachment is described by two parameters: effective division rate and effective detachment rate. We find that reversible attachment can be understood if we analyze behavior using lineage time (the time a lineage stays continually on the surface) rather than an experiment time, defined by time from inoculation. Specifically, reversible attachment comprises two regimes of behavior, defined by whether cells of the lineage stay on the surface long enough to divide, or not, before detaching. For lineages that detach before dividing at all, both PAO1 and PA14 behave similarly with near certain lineage “extinction,” wherein the entire lineage detaches. For lineages that stay long enough to divide, PAO1 and PA14 show surprisingly different behaviors. Our theoretical model provides a framework wherein time-dependent division and detachment rates and distributions of lineages can be extracted from our experiments. Our results suggest that PAO1 and PA14 utilize two fundamentally different surface colonization strategies. For PAO1, individual lineages commit relatively quickly to a surface compared to PA14, resulting in a steady progressive increase of a surface cell population that is irreversibly attached (i.e., committed to forming a biofilm). In contrast, PA14 lineages have high rates of cell detachment from surfaces. However, these detaching cells retain a memory of the surface (8), and ultimately form a planktonic population that is primed for attachment, so that sudden increases in irreversibly attached surface cell populations can occur. Our model provides a framework for categorizing different surface colonization strategies that lead to biofilm formation, and it is conceivable that in principle each approach has its own advantages under different circumstances.

## Results

### Two regimes of reversible attachment in PAO1 and PA14 are revealed through lineage analysis

When monitoring the number of cells on the surface as a function of the time from inoculation of the flow cell (denoted as experiment time), both strains follow a similar pattern (Figure 1). At early times, widespread detachment behavior is observed. Despite both division and additional attachment, the surface population essentially remains constant for a long and variable lag period (~10-20 h), after which the surface population will then begin to rise steeply, in a manner that can be fit to an exponential growth curve. However, further distinguishing their behaviors in a finer pitch of detail is difficult due to the random nature of reversible attachment. When comparing the surface population increases between PAO1 and PA14, we observe nearly all possibilities: we either observe that PAO1 has a steeper and earlier rise in the surface population compared to PA14, that PAO1 and PA14 have similar rises, or that PA14 has an earlier and steeper rise than PAO1 (Figure S1). Furthermore, it is difficult to correlate these observations with macroscopic crystal violet biofilm assays, where PAO1 has statistically significantly higher OD_550nm_ values compared to PA14 (Figure S2), which suggests that PAO1 is capable of forming early biofilms faster than PA14. PAO1 has a mean OD_550nm_ = 0.23 with a 95% confidence interval of (0.19, 0.26), while PA14 has a mean OD_550nm_ = 0.14 with a 95% confidence interval of (0.099, 0.18). Comparing the bootstrap sampling distributions of the mean OD_550nm_ values (which also generate the 95% confidence intervals) show that PAO1 has a higher mean OD_550nm_ value than PA14 (p-value of 0.0002). Using the median instead of the mean gives similar results. PAO1 has a median OD_550nm_ = 0.22 with a 95% confidence interval of (0.17, 0.29), while PA14 has a median OD_550nm_ = 0.12 with a 95% confidence interval of (0.094, 0.21). Comparing the bootstrap sampling distributions of the median OD_550nm_ values (which also generate the 95% confidence intervals) show that PAO1 has a higher median OD_550nm_ value than PA14 (p-value of 0.003). These apparently conflicting observations are not easily resolved with increased data collection since they arise from the intrinsic randomness of the process and not from incurring measurement errors. This instrinsic randomness, which is not uncommon in different aspects of biofilm formation, is usually neglected in analyses. In the present context, these effects complicate any analysis of the reversible attachment behaviors in PAO1 and PA14, that depend on traditional methods to monitor the number of surface cells as a function of experiment time or by macroscopic assays.

**Figure 1.**
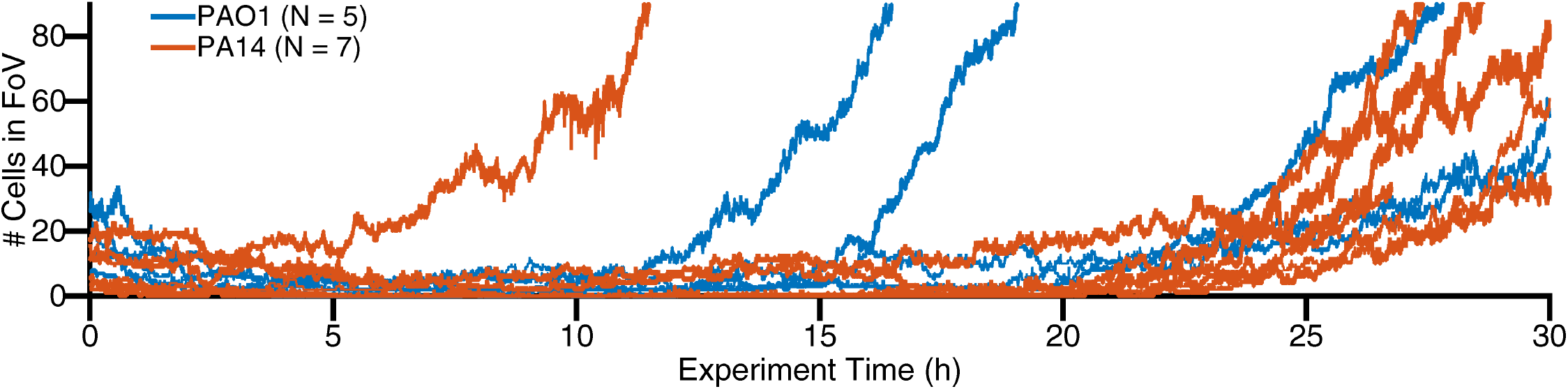
PAO1 and PA14 can both form biofilms and have similar trends of exponential surface population increase. Each line reprsents one experiment where we count how many cells are in a single field of view (FoV) for WT PAO1 and PA14 as experiment time progresses (5 and 7 independent experiments for PAO1 and PA14, respectively). Experiment time = 0 h corresponds to when imaging commenced after cells were inoculated into the flow cell chamber. Both PAO1 and PA14 have the variable lag period and the exponential increase, which is consistent with the fact that both strains initially undergo reversible attachment, and then subsequently form biofilms.

To account for the random nature of reversible attachment and the large fluctuations in the observations, we investigate the evolution of bacterial behavior as a function of surface sensing progression using lineage analysis. We monitor the time that a given isolated family, consisting of an attached cell (founder cell) and its progeny (daughter cells) via division, stays continually on the surface, which we designate as lineage time (*t* = *t*_lineage_). For each family, we begin tracking at the frame that an individual, founder bacterium attaches and assign this time as *t*_lineage_ = 0 h. We continue tracking until either the entire family detaches, or until we lose track of that family (where we can no longer distinguish individual cells, or the cells move out of the recording boundaries). This final time point is recorded as the family’s residence time. During reversible attachment regimes, families are categorized by whether a division event occurs or not before detaching. We denote families that detach before dividing at all as the “nonprocessive” regime of reversible attachment, and families that divide one or more times before detaching as the “processive” regime of reversible attachment, using language from enzyme kinetics. It is important to note that these regimes are distinct from irreversible attachment because during both regimes of reversible attachment, detachment is still prominent, while during irreversible attachment, detachment is much less common. All families analyzed here are shown in Figure 2 and Figure S3.

**Figure 2.**
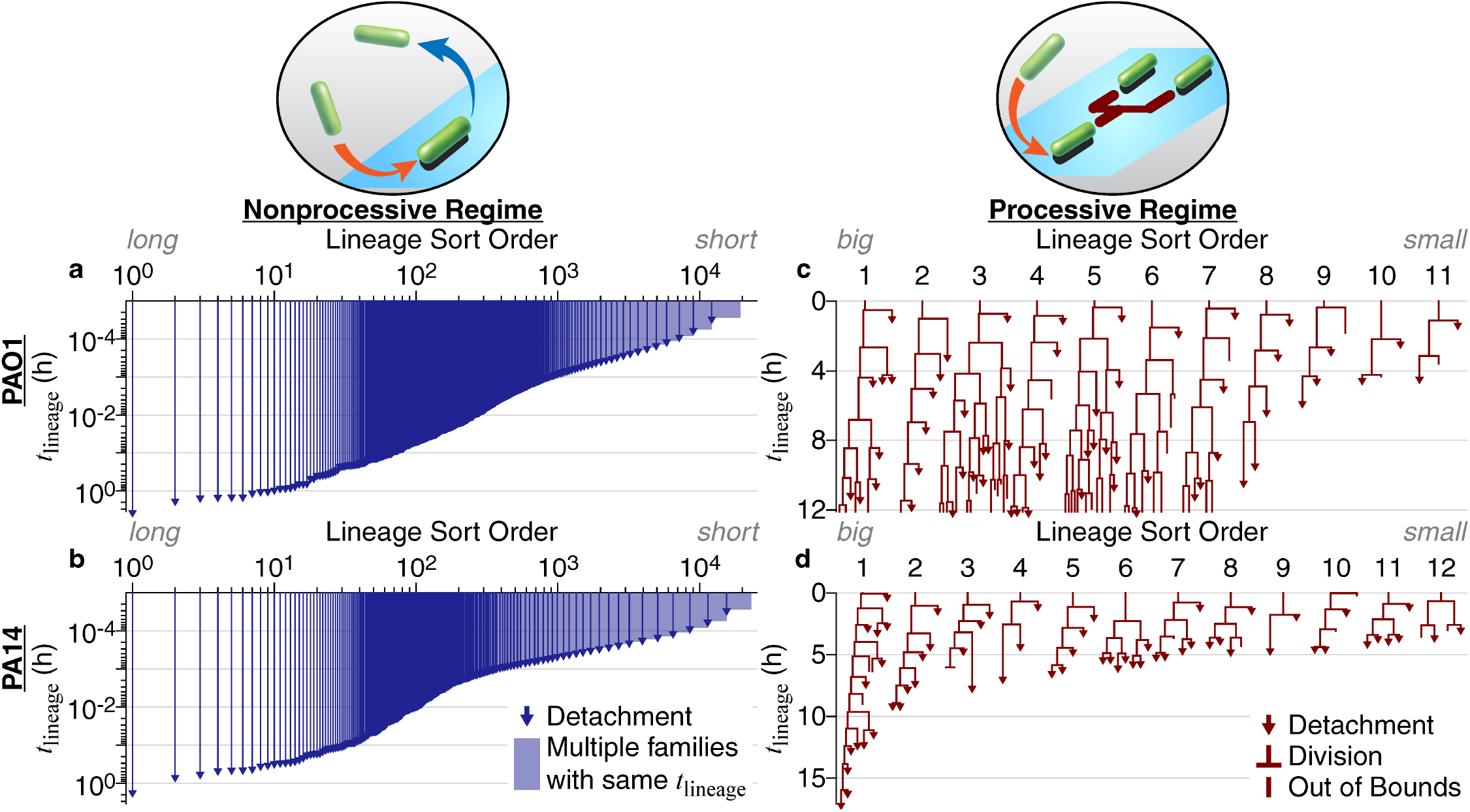
PAO1 and PA14 family trees in different regimes of reversible attachment. (a,b) Families in the nonprocessive regime of reversible attachment, which is when cells detach before dividing. Both axes are on a log scale. In this nonprocessive regime, PAO1 has *m* = 19353 tracked families, and PA14 has *m* = 23104 tracked families. Note the similarities between PAO1 and PA14. (c,d) Families in the processive regime of reversible attachment, which is when cells divide at least once before detaching. Both axes are on a linear scale. Each family start at *t*_lineage_ = 0 h when the founder cell attaches to the surface. Tracking continues for that family until either all members detach, or we lose track of the family (where we can no longer distinguish individual cells, or the cells move out of the recording boundaries). We then record this time as the family’s residence time. For each regime and strain, we sort families by residence times in descending order, which sorts them by the amount of time that they have continuously contacted the surface. In this processive regime, families here are used for the model and are a subset of the full tracked families shown in Figure S3.

Cells in both nonprocessive and processive regimes are present throughout the entire biofilm formation process. However, during the initial variable lag period, where the total surface population is not increasing, almost all cells are in the nonprocessive regime, while very few cells are in the processive regime. As experiment time elapses, the general observed trend is that cells in the processive regime become more common, while cells in the nonprocessive regime become less common, especially during period of surface population exponential increase. However, it is difficult to quantify such cellular behavior in this system because both regimes coexist with fluctuating proportions due to the inherent randomness in single cell behavior, thereby complicating any analysis of biofilm behavior as a function of experiment time. Thus, we utilize an analysis of lineage time to quantify the behavior of individual families in each regime.

When comparing the two regimes for either PAO1 or PA14, we find that the residence times are drastically different. In the nonprocessive regime, ~99% of cells stay on the surface for less than 30s for both strains. Furthermore, of the ~20,000 tracked families in the nonprocessive regime (both PAO1 and PA14), we observe less than 10 families (~0.05%) that have residence times comparable to the average doubling time of 1-2 h (Figure S4), which is the minimum residence time for families in the processive regime. Detachment dominates attachment and division in the nonprocessive regime, and essentially the surface population does not increase over the first 10-20 h of experiment time. In contrast, cells in the processive regime are in continuous contact with the surface for longer periods of time. Moreover, virtually all of the cells that remain surface engaged in the processive regime do so longer than cells in the nonprocessive regime. Finally, cells in the processive regime grow and divide on the surface and have clearly altered their behavior compared to “surface-naïve” planktonic cells, presumably as a consequence of activating surface sensing pathways.

Interesting trends emerge when comparing PAO1 and PA14 lineages in each regime. In the nonprocessive regime, we find that PAO1 and PA14 exhibit similar behaviors, where cells experience the surface transiently. However, in the processive regime, we see stark differences between PAO1 (44 families with 622 total descendants analyzed) and PA14 (31 families with 381 total descendants analyzed) (Figure S3). PAO1 families have more progeny retained on the surface, while PA14 families have more progeny detaching, which can be seen in a broad range of metrics. For example, we can compare single cell detachment behavior via the proportion of detachment vs division events. PAO1 has a statistically significantly lower proportion, with 143 (33%) detachment vs 289 (67%) division events, compared to PA14, with 130 (43%) detachment vs 175 (57%) division events, according to the *χ*^2^ test (p-value of 0.008). We can compare family-averaged detachment behavior with family tree asymmetry parameter Λ (8). Λ values closer to zero indicate a more symmetric family tree where more progeny are retained (more “two-legged” division nodes in the family tree, where both post-division daughter cells stay on the surface), while Λ values closer to one indicate a more asymmetric family tree where more progeny detach (more “one-legged” division nodes in the family tree, where one of the post-division daughter cells detach from the surface). PAO1 family trees have a median Λ = 0.33 with a 95% confidence interval of (0.25, 0.39), while PA14 family trees have a median Λ = 0.42 with a 95% confidence interval of (0.37, 0.52). Comparing the bootstrap sampling distributions of the median tree asymmetry values (which also generate the 95% confidence intervals) show that PAO1 family trees have a lower median Λ than PA14 family trees (p-value of 0.015). Overall, our data show that PAO1 and PA14 display similar behaviors during the nonprocessive regime, but during the processive regime, PAO1 shows a significantly higher likelihood of remaining surface-associated.

### “Divide-detach” stochastic model highlights differences between PAO1 and PA14 in the processive regime of reversible attachment

Our observations suggest that PAO1 is less prone to detachment than PA14. However, these metrics do not properly consider the collective time-dependent effects of division and detachment. For example, having more detachment events earlier in lineage time would have a much greater effect on the resulting family architecture compared to the same detachment events occurring several generations later. Even at the single cell level, gene expression is stochastic and can occur in a burst-like, intermittent manner (29), which contributes additional randomness to that cell’s behavior. Consequently, the behavior of an individual bacterium (in terms of whether or not in every instance they stay on the surface or detach after a division event) may be completely random and can only be described using statistical metrics. Since biofilm formation can be seen as an evolution of a population of random individual bacteria, it can be described as a stochastic process that depends on a number of control parameters as well as random environment variables. Consistent with that contention, large fluctuations are often observed in measured parameters (e.g., family trees), and these fluctuations are not easily mitigated with increased data collection since they arise from the intrinsic randomness of the process and not from incurring measurement errors. In general, although it is acknowledged that the unpredictability of single cell behavior can be important to surface sensing and biofilm development, this randomness is rarely accounted for in traditional microbiological studies.

To obtain more time-dependent comparisons that incorporate division and detachment effects, and to help account for the inherent randomness in observed family trees, we develop a “divide-detach” stochastic model. We use this model to study the temporal evolution of the expected number of surface cells in a family tree, or population size (30–35). In this model, the population size can increase or decrease by one bacterium as time evolves, and the population size can be infinite or null. The corresponding sample space Ω is given by Ω = {0, 1, 2, …, *N*, …}^*m*^, where *m* is the number of independent family trees, or different populations of bacteria. As time evolves, the population size can change and result in a sequence *ω* ∈ Ω, where *ω* is the set of family trees that are in the experiment. For example, if there is *m* = 1 family tree, then *ω* = {*ω*_1_}, and if there are *m* family trees, then *ω* = {*ω*_1_, …, *ω*_*m*_}. However, because this is a stochastic process, we cannot predict ahead of time what *ω* will be. Instead, what we know for *ω* are the set of possible observations (states) Σ and the actual observations *X*_*t*_ from experiments. The set of states is given by Σ = {0, 1, …, *N*, …}, where *N* represents the number of surface cells in a family and is infinite. Σ tells us what observations (number of surface cells) are possible for any family tree during an experiment. The actual observations of *ω* are given by 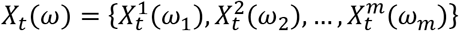, which is how many surface cells are observed in each of the *m* family trees at time point *t* during an experiment, and *X*_*t*_(*ω*) is a random variable *X*_*t*_: *ω* → Σ that defines this stochastic process. Having a random variable means that for the family trees *ω* and each time point *t*, we observe *X*_*t*_(*ω*) taken from the set of states Σ according to a certain (not necessarily known) probability distribution; but when we repeat the experiment, we will not necessarily observe the same *X*_*t*_(*ω*) for the same time point *t* and family trees *ω*. Figure 3 shows examples of this process for *m* = 1 family tree (Figure 3a) and for *m* = 3 family trees (Figure 3b).

**Figure 3.**
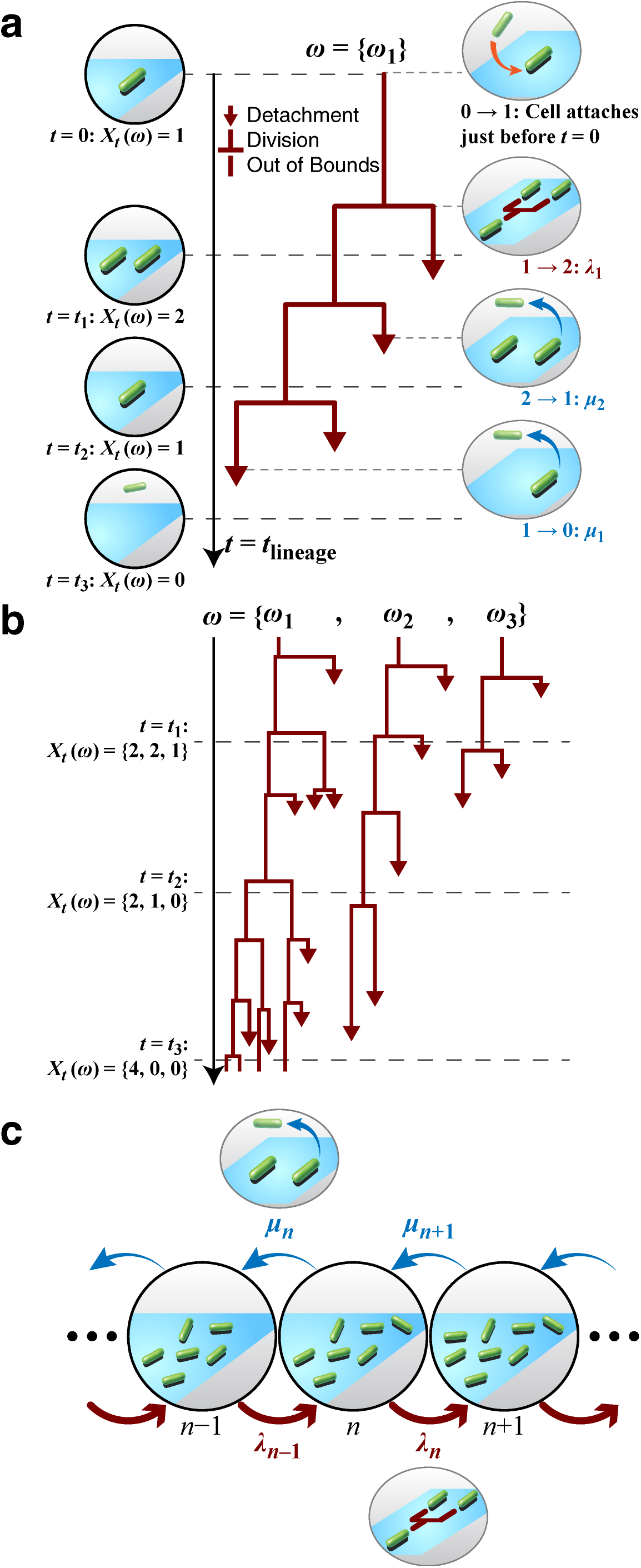
Family trees are a stochastic process. (a) Example illustrating the stochastic process with *m* = 1 family tree. In this case, at time *t*, *X*_*t*_(*ω*) is the number of observed cells in the family tree *ω* = {*ω*_1_} at time *t*. Attachment of the founder cell happens just before time *t* = *t*_lineage_ = 0, so it is not explicitly captured by this process. When a cell divides, it undergoes a transition *n* → *n* + 1 = *λ*_*n*_, and when it detaches, it undergoes a transition *n* → *n* − 1 = *μ*_*n*_, where *n* is the state (i.e., number of observed cells) before the transition. (b) Example illustrating the stochastic process with *m* = 3 family trees. At time 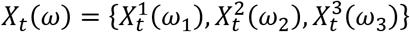 are the number of observed cells for each of the family trees *ω* = {*ω*_1_, *ω*_2_, *ω*_3_}. (c) Dynamics of the stochastic process for state *n*. As shown in part (a), a transition *n* → *n* + 1 = *λ*_*n*_ occurs when a cell divides, and a transition *n* → *n* − 1 = *μ*_*n*_ occurs when a cell detaches.

The dynamics of such a stochastic process are given by the evolution of the probability distribution *P*(*j*, *s* + *t*|*i*, *s*), which gives the probability of transitions between all states for all *t* ≥ 0 and can be rewritten as *P*_*t*_(*j*|*i*). For a family tree, the only possible transitions are the neighboring transitions, *n* → *n* + 1 = (*n* + 1|*n*) = *λ*_*n*_ and *n* → *n* − 1 = (*n* − 1|*n*) = *μ*_*n*_. As a result, the dynamics of this process can be described by looking only at the evolution of the probability distribution *P*_*t*_(*n*|*n*) for state *n* ∈ Σ. The rates *λ*_*n*_ and *μ*_*n*_ determine the intensity of increase (i.e., division) or decrease (i.e., detachment), respectively, for state *n*. In a family tree, each cell can divide (with a division rate *λ*) or detach (with a detachment rate *μ*), so the rates become *λ*_*n*_ = *λn* and *μ*_*n*_ = *μn*. Figure 3c shows a schematic of the dynamics described here (i.e., how the population size can increase or decrease).

The equation describing the evolution of this process is given by the Kolmogorov backward equation, also called the master equation, which reads

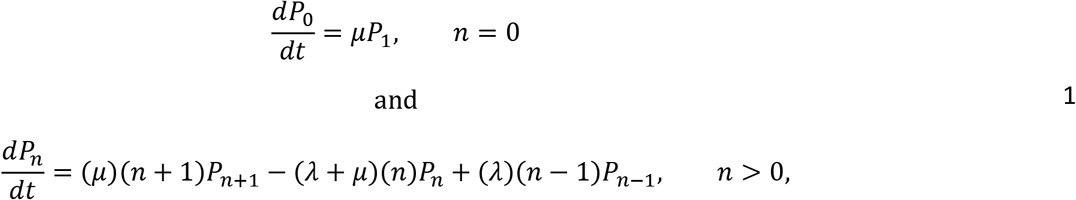

where *P*_*n*_ = *P*_*n*_(*t*) = *P*_*t*_(*n*|*n*), *P*_*n*+1_ = *P*_*n*+1_(*t*) = *P*_*t*_(*n* + 1|*n*), and *P*_*n*−1_ = *P*_*n*−1_(*t*) = *P*_*t*_(*n* − 1|*n*). We refer the readers to the methods to find the details of the solution to this equation.

Experimentally, by having access to *m* independent family trees, it is possible to build the probability distribution by counting the number of families that have zero cells, one cell, two cells, and so on, at a given lineage time *t*. In other words, for each time point *t*, we plot the actual observations *X*_*t*_(*ω*) on a histogram to derive the probability of each of the states *n* occurring. For families in the processive regime of reversible attachment, we avoid potential problems arising from tracking limitations by selecting *m* = 11 families for PAO1 (out of 44 families) and *m* = 12 families for PA14 (out of 31 families), with a *t*_lineage_ range of 0-12 h for PAO1 and 0-10 h for PA14 (see Figure S3 caption for family selection criteria).

Comparing the experimental data with the model is not straightforward when using the probability distributions directly. In the experimental data, there are a finite number of families and a finite number of cells in a family, which means that it is difficult to generate distributions that are well populated for quantitative comparisons. To overcome this limitation, we employ the method of moments, which provides information about the distributions, to fit the model to experimental data and obtain the rates. Instead of comparing the experimental and model probability distributions *P*_*n*_(*t*), we compare the experimental and model moments, 〈*n*(*t*)^*k*^〉, where *k* is the k-th moment. We can calculate the experimental moments directly from the experimental probability distribution, and we can obtain the model moments from the model probability distribution (eq. 2) given by the master equation (eq. 1). The equations for the model moments are shown in the methods (eq. 3-4). To compare experiment with model, we use the first two moments. The first moment is the mean, and the second moment is related to the variance, since the variance equals the second moment minus the first moment squared.

When we plot the moments calculated from the experimental data for families in the processive regime of reversible attachment (Figure 4), striking differences between PAO1 and PA14 are revealed. PAO1 follows an exponential growth curve, while PA14 follows a Gaussian curve. These curves are consistent with what we see in the family trees. For PAO1, many of the families have increasing number of cells, while for PA14, fewer of these families are present, and most families end in detachment. However, as we have previously shown, PA14 cells that detach have already initiated the surface sensing process, and they retain memory of the surface based on their prior surface residence, which primes them for subsequent irreversible attachment (8). Also, from our data, the variances for both PAO1 and PA14 can be as large as the mean population size, indicating that extinction in an individual lineage can happen at any time, even in a population that is exponentially growing on average. Therefore, it is important to note that individual lineage “extinction” events (where the entire family detaches) do not indicate a failure to form a biofilm.

**Figure 4.**
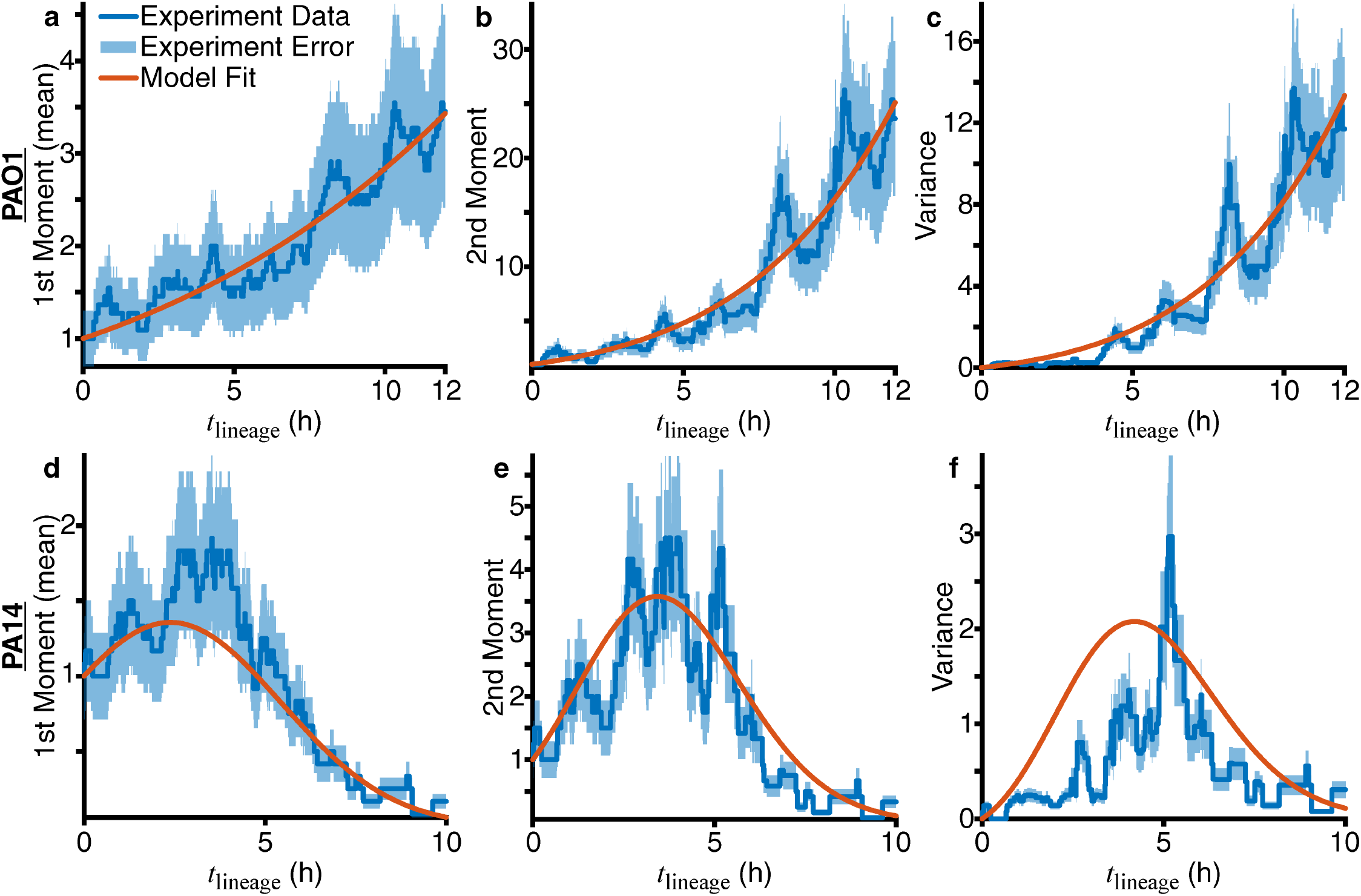
Obtaining division (*λ*) and detachment (*μ*) rates by fitting experimental and model moments of the number of cells in a family for families in the processive regime of reversible attachment. Moments and variance calculated from experimental data are plotted as blue lines, with the relative error (calculated as 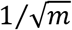, where *m* is the number of families used) shown as the light blue shaded area. Variance is defined as the second moment minus the first moment squared. Red lines show the fits to the first and second moments (eq. 3-4) using nonlinear least-squares. For the model, we use the linear functional form of the rates, *λ*(*t*) = *L*_0_ + *L*_1_*t* and *μ*(*t*) = *C*_0_ + *C*_1_*t*. The resulting coefficients from the fits are as follows: for PAO1, *L*_0_ = 0.136 h^−1^, *L*_1_ = 0 h^−2^, *C*_0_ = 0.0242 h^−1^, *C*_1_ = 0.00147 h^−2^, and for PA14, *L*_0_ = 0.256 h^−1^, *L*_1_ = 0 h^−2^, *C*_0_ = 0 h^−1^, *C*_1_ = 0.107 h^−2^.

With our model, the temporal evolution of a family tree can be described by the single cell division (*λ*) and detachment (*μ*) rates. *λ* is likely related to cellular events that contribute to surface growth, which can be affected by complex factors such as changes in cellular metabolism or the local availability of nutrients. Likewise, *μ* is likely to be related to cellular events that contribute to detachment, such as the production of EPS and the activities of motility appendages. Both rates can be time-dependent in principle, so *λ* = *λ*(*t*) and *μ* = *μ*(*t*). For example, as bacteria continue proliferating on the surface, they can spend more of their metabolic energy towards EPS production rather than for division, and they can start detaching less. However, finding the exact functional form of time dependence to use in the model is difficult. We first start with the simplest form of time-dependence (linear, or first order polynomial), where *λ*(*t*) = *L*_0_ + *L*_1_*t*, *μ*(*t*) = *C*_0_ + *C*_1_*t*, and {*L*_0_, *L*_1_, *C*_0_, *C*_1_} are the coefficients that we obtain by fitting the experimental data to the model. *λ*(*t*) and *μ*(*t*) are rates that represent probabilities per time unit, which means they are positive and have dimensions of inverse time, [*λ*] = [*μ*] = [time]^−1^. Therefore, the coefficients *L*_0_ and *C*_0_ also have dimensions [*L*_0_] = [*C*_0_] = [time]^−1^, and the coefficients *L*_1_ and *C*_1_ have dimensions [*L*_1_] = [*C*_1_] = [time]^−2^. By dimensional analysis, we can extract time scales for lineage-level growth (via division) and death (via detachment) behaviors from either the rates (*λ*^−1^ and *μ*^−1^) and the coefficients (*L*_0_/*L*_1_ and *C*_0_/*C*_1_). If the experimental data and model do not show good agreement, then we can reiterate this process with progressively more complicated functions. Additionally, the shape of the experimental moments can guide us in choosing the correct function for the rates.

With linear time dependence, we already obtain good agreement when fitting using nonlinear least-squares, as shown by the results of the model fits to the experimental moments in Figure 4. To ensure the fit results give meaningful coefficient values, we set the following constraints based on experimental data. The rates are positive, so *λ*(*t*) > 0 and *μ*(*t*) > 0. As seen in the family trees in Figure 2 and Figure S3, division events are roughly evenly spaced out in time, and cells are not nutrient-limited inside the experimental system, so *λ*(*t*) should be constant. Thus, we set *L*_1_ = 0, and *λ*(*t*) = *L*_0_. We consider any coefficient < 10^−5^ as zero for subsequent analysis based on the precision of the experimental data. The resulting coefficients from the fits are as follows: for PAO1, *L*_0_ = 0.136 h^−1^, *L*_1_ = 0 h^−2^, *C*_0_ = 0.0242 h^−1^, *C*_1_ = 0.00147 h^−2^, and for PA14, *L*_0_ = 0.256 h^−1^, *L*_1_ = 0 h^−2^, *C*_0_ = 0 h^−1^, *C*_1_ = 0.107 h^−2^.

We find that *μ*(*t*) is time-dependent for both strains (i.e., *C*_1_ is non-zero). For PAO1, we find that *μ* is slowly increasing, since *C*_1_ is ~1 order of magnitude smaller than *C*_0_, and both coefficients are positive and smaller than *L*_0_. For PA14, *μ* is increasing quite rapidly, since *C*_1_ is positive and is much greater than *C*_0_. Because *μ*(*t*) is time-dependent, the relevant time scale *τ*_*μ*_ to extract for time-dependent lineage-level detachment behavior for both PAO1 and PA14 is *τ*_*μ*_ = *C*_0_/*C*_1_ (≈16 h for PAO1, and 0 h for PA14). Also, because *λ*(*t*) is constant, the relevant time scale *τ*_*λ*_ to extract for time-independent lineage-level division behavior for both PAO1 and PA14 is 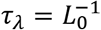 (≈7 h for PAO1, and ≈4 h for PA14). These values are consistent with the experimental data. Interestingly, for both strains, we find that *τ*_*λ*_ is bigger than the average division time by a factor of ~3, which means that *τ*_*λ*_ corresponds to ~3 generations of division. The value of *τ*_*λ*_ corresponds closely to the time that a given lineage persists on the surface before ultimately going extinct and detaching (i.e., residence time). To calculate the mean residence times of the experimental lineages in Figure 2, we only include lineages that ultimately detach before the cutoff time (12 h for PAO1, 10 h for PA14). This results in ≈6 h for PAO1 and ≈5 h for PA14, which are very close to the *τ*_*λ*_ values obtained from the coefficients (≈7 h for PAO1, and ≈4 h for PA14, see above). For PAO1, having a larger *τ*_*λ*_ and a slowly increasing and relatively small *μ* mean that lineages are division-dominant (*λ* > *μ*) as they spend more time on the surface. Rather than ultimately detaching, we see many lineages persist on the surface and increase their number of cells despite having detachment events. At *t*_lineage_ = 12 h, 7 of the 11 families still exist on the surface (Figure 2c, #1-7). For PA14, having a smaller *τ*_*λ*_ and a rapidly increasing *μ* means that lineages are initially division-dominant (*λ* > *μ*), but then become detachment-dominant (*μ* > *λ*) after a certain amount of time on the surface, which is also the time scale described by *τ*_*λ*_. We see that many lineages grow to at least 2-3 generations, which corresponds to the lineage time where *λ* > *μ*. Once *μ* > *λ*, then many families begin to detach until, at *t*_lineage_ = 10 h, only 1 of the 12 families still exists on the surface (Figure 2d, #1). Clearly, unlike PAO1, PA14 cells that have started the surface sensing process do not necessarily stay on the surface. Rather, as we have shown previously (8), they rejoin the planktonic population as “surface-sentient” cells that are primed for longer surface residence times during subsequent attachment. Similarly, the value of *τ*_*μ*_ corresponds closely to cellular activities that affect detachment, such as, for example, the competition between EPS production and motility appendage activity. EPS is likely to affect detachment more for PAO1 than for PA14, since PAO1 is known to produce the Psl EPS, while PA14 cannot. On the other hand, presumably because of the Pil-Chp system, motility appendage activity is likely to affect detachment more for PA14 than for PAO1. Consistent with this hypothesis, PAO1 has a much larger *τ*_*μ*_ compared to PA14 (which is zero).

With these model parameters, we can evaluate the model probability distribution *P*_*n*_(*t*) to then compare with the experimental probability distribution *P*_*n*_(*t*). We show the comparisons of the probability distributions with two different visual representations in Figure 5. The first is plotting *P*_*n*_(*t*) vs *n* for specific lineage times *t* = {2.5, 5, 7.5, 10} h, and the second is plotting the entire *P*_*n*_(*t*) vs *n* and *t* as a contour plot. The plots of the probability distributions also show good agreement. For the probability contour plots, agreement between experiment and model are assessed as follows. The regions of high (*P*_*n*_(*t*) ~ 1) and low (*P*_*n*_(*t*) ≤ 10^−2)^ probability contours are similar in shape and location (in the *n*, *t* plot space) between experiment and model. The shapes of the probability contours are consistent with the plots of the moments in Figure 4. For PAO1, as *t* progresses, the probability of having more cells per family (higher *n*) increases. For PA14, the probability of having higher *n* increases and then decreases as *t* progresses. However, further direct comparisons of the probability distributions are difficult. As previously mentioned, the experimental probability distributions will invariably be sparser than the model probability distributions, which can be seen in the plots as either jagged lines or holes in the contours. This sparseness comes from having finite experimental data and is not straightforward to remove (e.g., via interpolation). Nevertheless, the model probability distribution can be used to describe what family tree architectures we expect to observe during similar experiments.

**Figure 5.**
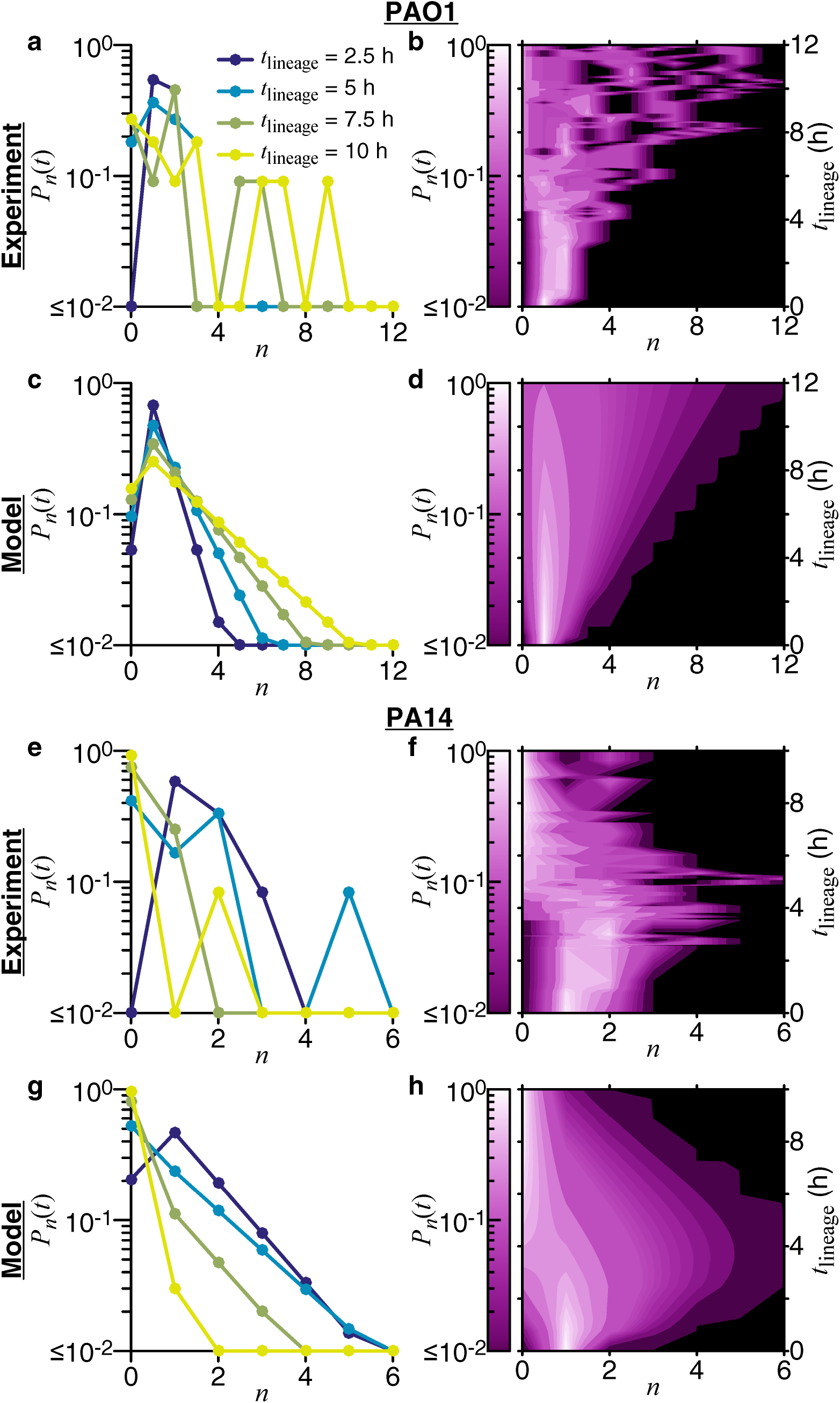
Comparing experimental and model probability distributions for the number of cells in a family for families in the processive regime of reversible attachment. Experimental probability distributions are built directly from the data, as described in the methods (section “Multigenerational family tracking analysis”). For the model probability distribution *P*_*n*_(*t*), we use eq. 2 and the linear functional form of the rates, *λ*(*t*) = *L*_0_ + *L*_1_*t* and *μ*(*t*) = *C*_0_ + *C*_1_*t*. The model rate coefficients used are shown in Figure 4. Probability distributions are compared in two different ways. The left column shows plots of *P*_*n*_(*t*) vs *n* for specific lineage times *t* = {2.5, 5, 7.5, 10} h, and the right column shows plots of the entire *P*_*n*_(*t*) vs *n* and *t*, where the probability is represented by the shades of color in the contour plots. Probabilities are shown on a log scale.

The model can also be applied for cells in the nonprocessive regime of reversible attachment. We find that for both PAO1 and PA14, the moments fit to an exponential decay function (Figure 6). This is what the model predicts if there is only detachment and no division, and it correctly describes the data, because cells in the nonprocessive regime detach before dividing on the surface. Furthermore, the variances are of the same order of magnitude as the mean population, which means that lineages can become extinct at any time. Thus, for both processive and nonprocessive regimes of reversible attachment, the stochastic model described here accurately describes the behaviors of PAO1 and PA14, including their differential paths to irreversible attachment.

**Figure 6.**
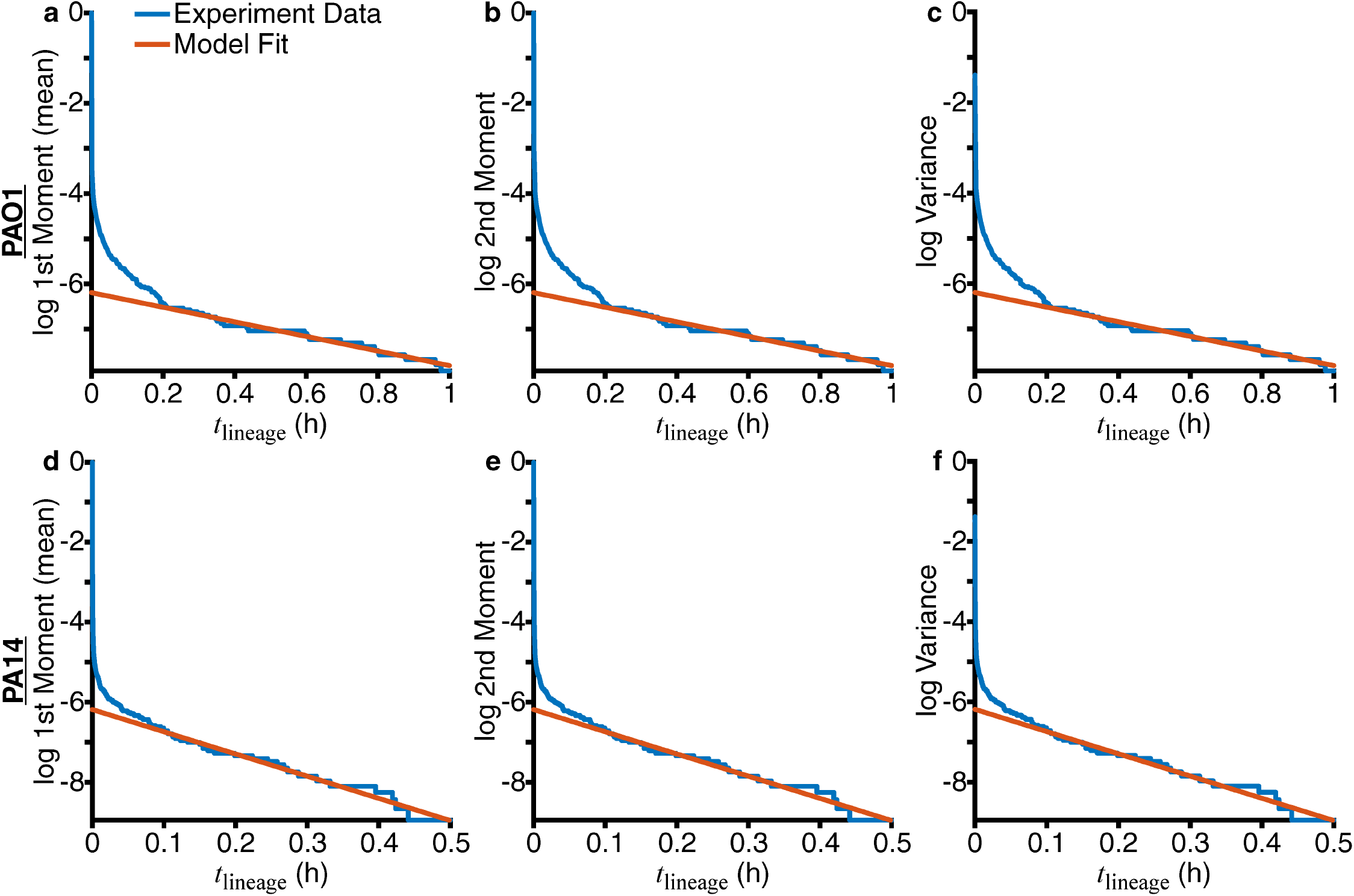
Fitting experimental moments show that families in the nonprocessive regime of reversible attachment have only detachment events. The natural logarithm (log) of the moments and variance calculated from experimental data are plotted as blue lines. Relative errors (calculated as 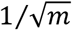, where *m* is the number of families used) are not plotted here, as *m* ~ 20000 for both PAO1 and PA14, and the values are very small. Red lines show the fits of the experimental data to the function log(*y*(*t*)) = *a*_1_*t* + *a*_0_. For PAO1, fits were performed for 0.2 h ≤ *t* ≤ 1 h, with the resulting fit coefficients as *a*_1_ = −1.61 h^−1^, *a*_0_ = −6.19. For PA14, fits were performed for 0.1 h ≤ *t* ≤ 0.5 h, with the resulting fit coefficients as *a*_1_ = −5.55 h^−1^, *a*_0_ = −6.19.

### PAO1 and PA14 have distinct progressions of surface colonization, which suggest contrasting surface engagement strategies

Our observations and results imply that both PAO1 and PA14 start their initial surface engagement with similar behavior (in terms of surface residence times) in the nonprocessive regime of reversible attachment, but then they diverge strongly in the processive regime of reversible attachment. PAO1 shows an increase in the number of families that commit relatively quickly to surface growth, and this is likely a factor that contributes to the trend of PAO1 forming faster biofilms compared to PA14 as seen in the crystal violet assays (Figure S2). For PAO1, this early attachment behavior can be more intuitive when correlating with the general progression of biofilm formation. PA14, in contrast, shows a larger number of detachment-dominated families even though the entire population eventually forms a biofilm, which can be a counterintuitive result. Compared to PAO1, where production of sticky EPS appears to be the dominant mechanism driving irreversible attachment (22, 23), PA14 appears to utilize a different surface colonization strategy dependent on progressive suppression of surface motility appendage activity (8), but it is not obvious is how this motility suppression strategy can lead to rapid changes in bacteria detachment rates from surfaces, as shown by the model.

We investigated how detachment events occur for PA14 to gain insight into this alternate surface colonization strategy and why appendages and their activities can give rise to a time-dependent detachment rate *μ*. Consistent with previous results where flagellum-mediated surface spinning generally results in a detachment event (36), we find that ~90% of detachment events occur when a cell has the mature flagellum inherited from its ancestor, as opposed to that cell having to form a new flagellum post-division (Figure 7a,c). Interestingly, deleting the *pilA* gene (Δ*pilA*, missing the major subunit of the TFP filament) results in significantly fewer detachment events (*χ*^2^ test p-value≪ 10^−4^) for cells that have a mature flagellum. Compared to WT, only roughly half of detachment events occur when the cell has a mature flagellum in the Δ*pilA* mutant (Figure 7b,c), an observation that suggests that TFP are important to the detachment process. For the Δ*pilA* mutant (and to a much lesser extent in WT), we also observe detachment events with cells that did not have a labeled flagellum, which suggests that non-flagellum-mediated detachment events can also occur. To study how TFP can influence flagellum-mediated spinning and detachment, we adapt a previously developed hydrodynamic model (37). Simulations show that TFP activity (i.e., extension or retraction) can lead to changes in the cell body tilt angle relative to the surface. In the case where the non-flagellated pole is attached to the surface, TFP extension during flagellum-mediated spinning results in the cell tilting to near vertical orientations, while retraction results in a smaller tilt angle (Figure 7d). During flagellum-mediated spinning, near vertical orientations correlate with higher rates of detachment, while orientations closer to the surface correlate with a decreased likelihood of detachment (36, 38). Consistent with previous results, the cell without TFP is more likely to assume an orientation closer to the surface (i.e., horizontal), while the cell with TFP extended the entire time is more likely to assume a near vertical orientation (8).

**Figure 7.**
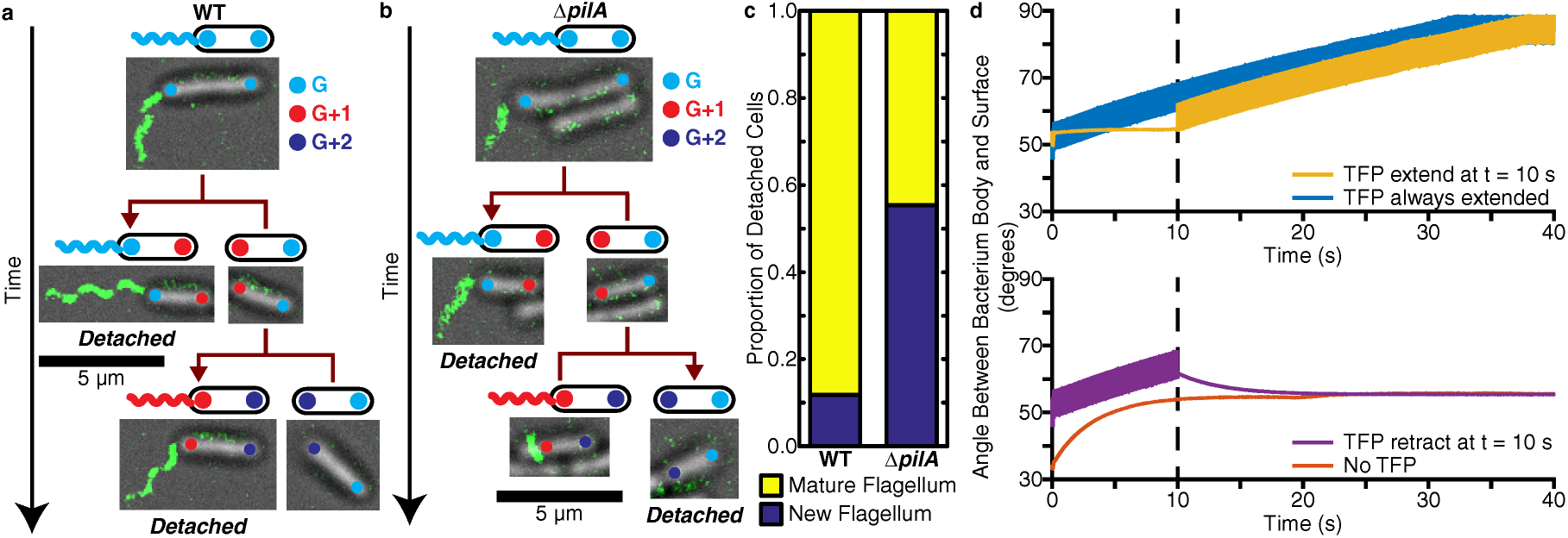
TFP and flagella are both important for the detachment process in PA14. (a,b) The location of the flagellum and the age of poles (measured in generations) can be tracked across multiple generations. The fluorescence image of the labeled flagellum is overlaid on top of the bright field image, and the poles are overlaid as colored circles (color representing the pole generation *G*). Scale bars for the pictures are 5 μm. (a) Example tracking for WT, where cells with mature flagella detach. (b) Example tracking for the Δ*pilA* mutant (deletion of the major subunit for the TFP filament), where one cell with a mature flagellum and one cell without a flagellum detach. (c) Proportion of detached cells with a mature flagellum vs a new flagellum (which includes no flagellum), calculated from 154 events for WT and 74 events for Δ*pilA*. The proportions are statistically significantly different between the strains according to the *χ*^2^ test (p-value≪ 10^−4^). (d) Angle that the bacterium’s body makes with the surface for different TFP conditions in the hydrodynamic model: TFP extension at t = 10 s (top plot, yellow-orange), TFP retraction at t = 10 s (bottom plot, purple), TFP extended throughout (top plot, blue), and no TFP (bottom plot, red). If the bacterium does not spin, then the angle between the body and surface will stay at the (arbitrary) initial condition we have chosen in the model. We show time using units of seconds and a torque value of 2 pN μm (37).

These results suggest that detachment rates are higher when TFP activity and flagellum activity are high and/or coincide, and that detachment rates are lower when the activities are reduced and/or do not coincide. Given that PA14 has small average family size, small surface residence times, and large surface detachment rates, observations of suppression of both appendage activity and detachment are expected to be extremely rare during reversible attachment (i.e., while cells are transiently on the surface). Nevertheless, in our family tree data, we can find examples where we can compare cells from the same generation but on different branches of the family tree. In these cases, we observe detachment in branches where appendage activities are high (and/or coincide), and no detachment in branches where appendage activities are reduced and/or do not coincide (Figure 8). In example (i), we see that appendage activity is reduced around *t*_lineage_ ~ 6 h, which coincides with the presence of a division event where no daughter cells detach. In examples (ii) and (iii), appendage activity does not become quiescent and detachment continues to occur for subsequent division events. This appendage activity analysis was repeated with Δ*pilA* for validation and was consistent with previous results (Figure S5).

**Figure 8.**
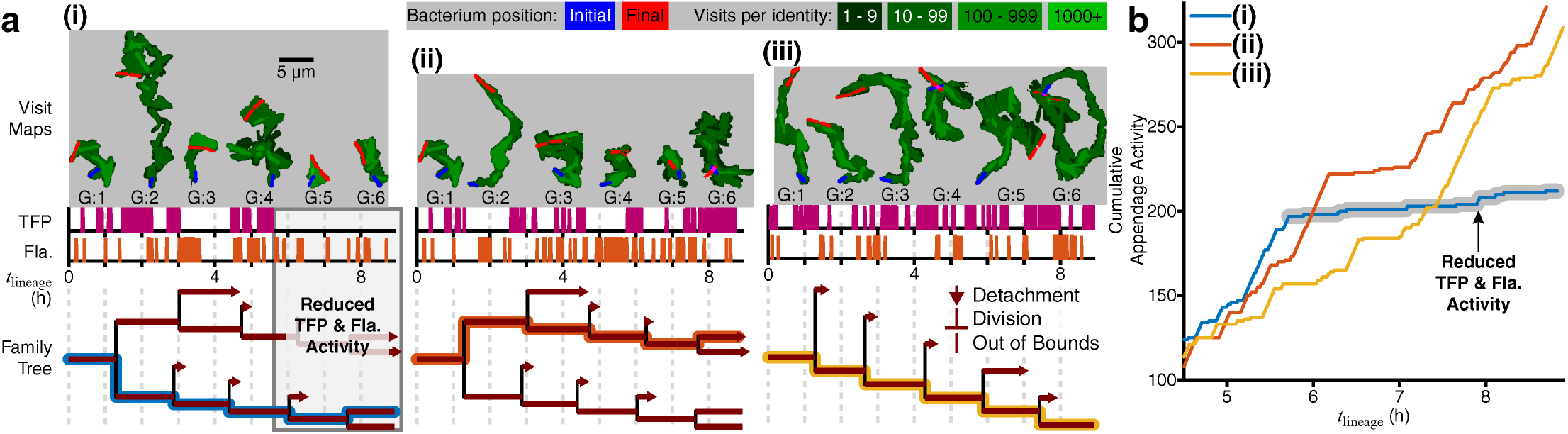
Family tree architecture controlled by generation-dependent motility activity. (a) TFP and flagellum activities are plotted for single branches of a family as magenta and red spike plots, with each spike representing one instance of activity. The colored line on the family tree plot traces the analyzed branch. We plot the corresponding visit map of each single cell (top, gray boxes), shown as shades of green in a logarithmic color scale with initial (final) positions shown in blue (red), to show the trajectory history and visual representation of TFP and flagellum activity. The size of each visit map is roughly proportional to the level of activity: cells with high TFP activity have elongated visit maps, while cells with high flagellum activity have circular visit maps. For WT, we see one example where TFP and flagellum activity are reduced and do not coincide, resulting in a division event where no daughter cells detach (i). The other examples show either higher activities or activities that coincide, resulting in a continuation of division events with detachment occurring (ii, iii). (b) Plot of the cumulative appendage activity (i.e., the cumulative number of instances of either TFP or flagellum activity) for the 3 examples in (a). For example (i), the curve plateaus out at *t*_lineage_ ~ 6 h, which is when the reduction in appendage activity occurs. For examples (ii) and (iii), the curves continue to rise, which shows that the appendages are continuing to be active.

## Discussion

Clearly, the application of stochastic models can be quite powerful in understanding microbiological systems that involve strong fluctuations. The behavior of each lineage is a record of how a specific cell and its progeny managed to stay and proliferate on the surface during cellular changes induced by surface sensing, which has multigenerational consequences. Even though the probability of a specific cell attaching to a surface and proliferating successfully is initially vanishingly small, surface sensing can modify outcomes by changing the structure of family trees, as we can see from the evolution of reversible attachment from the nonprocessive to processive regimes, for example. Interestingly, that the process of reversible attachment can be described by a stochastic model is telling: whether a bacterium encountering a surface makes it to irreversible attachment and eventually participates in biofilm formation may be quantitatively cognate to the description of whether patient zero’s disease will die out after a few infections, or take hold and become an epidemic. The fact that biofilm formation seems to inevitably happen is due to factors such as the large number of lineages that encounter the surface, and the existence of multigenerational memory, which can mitigate against initial failure to attach by conditioning a planktonic population primed for improved subsequent attachment.

Indeed, a recent study applied a variation of our approach to antibiotic treatment of bacteria (39). In fact, the quantitative evolution of bacterial populations in early biofilm formation is analogous to a time-reversed version of antibiotic treatment: the nonprocessive regime of reversible attachment behaves like bacterial population dynamics for antibiotic treatment well above the minimum inhibitory concentration (MIC). In the present study, however, we are able to perform an unprecedented level of longitudinal comparison between theory and experiment. Because we have information on the fates for every cell in a large number of bacterial lineages that occur during early biofilm formation, we can directly measure and analyze the time evolution of the system. This analysis provides a conceptual framework for understanding the taxonomy of surface colonization strategies and reveals an unanticipated difference between PAO1 and PA14 behavior.

One of the old questions about biofilm formation is whether it is the newly landed cells or the dividing cells on the surface that contribute more to the biomass increase in the biofilm. Our results suggest that not only is the answer species and strain dependent, the question is misleading because of the assumed either-or format of the answer. Surface sensing can evolve progenitor cells which land on a surface and commit almost its entire division lineage to the surface, thereby drastically increase biomass.

### Complementary surface colonization strategies: immediate vs deferred gratification

The “divide-detach” stochastic model highlights two distinct but complementary strategies for surface colonization that are illustrated by PAO1 and PA14. For PAO1, surface population increase takes the form of the few families that are more successful in retaining surface progeny. PAO1 families generally stay on the surface during biofilm formation, likely due to the Wsp surface sensing system and Psl EPS secretion. Previous work has shown that early surface attachment behavior depends on EPS production via the Wsp system (9, 23). In contrast, for PA14, surface population increase takes the form of many families that are less successful in retaining surface progeny due to surface detachment. However, PA14 cells can “remember” the surface due to the Pil-Chp system and multigenerational cAMP-TFP memory (8), which primes them for biofilm formation whether they are currently on the surface or not and eventually leads to progressive suppression of motility appendage activity. Both strategies are viable for surface colonization. PAO1 cells tend to attach, increase their surface population more quickly, and persist longer on a surface compared to PA14, which suggests that PAO1 can potentially attach to surfaces even in ecologically crowded environments or successfully form biofilms by outgrowing competing species. Indeed, this has been experimentally observed: EPS-producing *P. aeruginosa* strains tend to persist on surfaces better than EPS nonproducers, despite possible exploitation by “cheaters” that can potentially use the communal good of EPS (40). In contrast, PA14 exposed to a surface do not initially stay on the surface, and slowly increase surface coverage. Rather, they form a surface-sentient planktonic population that can quickly attach and colonize the surface later in time, which may be better adapted for overwhelming host defense (i.e., a naïve surface) rather than microbial competition. Moreover, it is interesting to note that EPS secretion is extracellular and can be shared spatially and temporally with both neighbors from different lineages and descendants in close proximity (41), whereas memory is intracellular and can be only passed down temporally through division.

It is possible that our observations and results with PAO1 and PA14 may be generalizable to other *P. aeruginosa* strains. The majority of strains in the International *Pseudomonas* Consortium Database (IPCD) can be identified as either PAO1-like or PA14-like based on their phylogeny (i.e., same phylogenetic sub-group as either PAO1 or PA14) (42–45). Consistent with our results, crystal violet biofilm assays show that the PAO1-like strains seem to produce early biofilms faster than the PA14-like strains (Figure S6). Although it is clear from the data spread that there is more to *Pseudomonad* phylogenetic diversity than biofilm behavior, this observation suggests that the phylogenetic distance from either PAO1 or PA14 could be incorporated into a metric for categorizing a *P. aeruginosa* strain’s biofilm formation behavior as either PAO1-like or PA14-like. It is tempting to draw an analogy between differences in these strategies to differences in “immediate-” vs “deferred-gratification” behavior in a prototypical cognitive-affective processing system, with the latter correlating to successful outcomes in complex competitions (46). For *P. aeruginosa*, there is no cognition of course, but the existence of a specific sensing cascade for PA14 effectively encodes the analog of “deferred-gratification” behavior at a molecular level. Indeed, PA14 is usually considered to be more virulent than PAO1 (47). However, it is likely that these bacterial strategies have their own advantages under different circumstances. Furthermore, our model can be applied to other bacterial systems to understand how they utilize their cellular machinery for various surface colonization strategies.

## Materials and Methods

### Strains and growth conditions

*Pseudomonas aeruginosa* PAO1 and PA14 wild type (WT) strains were used in this study. For the flagellum localization data, PA14 WT and Δ*pilA* (deleting the major subunit of the TFP filament) (48) with FliC (the major subunit of the flagellum filament) modified to FliC(T394C) (49) were used. PAO1 was cultured as previously described (21, 23), and PA14 was cultured as previously described (8). Culturing protocols are summarized as follows. Bacteria were plated on LB agar plates and incubated at 37 °C overnight. Individual colonies were swabbed from the plate and grown overnight for ~18 h in an incubator at 37 °C shaking at 220 rpm. Overnight cultures were regrown in the same overnight growth conditions to an OD_600nm_ ~ 0.4-0.6. Regrowth cultures were then diluted in flow cell conditions to an OD_600nm_ ~ 0.01-0.03. These final diluted cultures were used for injection into the flow chamber.

Different medium conditions were chosen for PAO1 and PA14 based on the medium optimized for flow cell early biofilm formation experiments for each individual strain in prior work. For PAO1, overnight and regrowth media consisted of FAB medium with 30 mM glutamate, while flow cell media consisted of FAB medium with 0.6 mM glutamate (21, 23). For PA14, overnight and regrowth media consisted of M63 medium with 1 mM magnesium sulfate, 0.2% glucose, and 0.5% casamino acids (CAA), while flow cell media consisted of M63 medium with 1 mM magnesium sulfate, 0.05% glucose, and 0.125% CAA (8, 48). For flagellum staining experiments, the flow cell media also contained 0.375 μg/mL Alexa Fluor 488 C5 maleimide dye (Molecular Probes). For more details on the culturing procedures, please refer to the corresponding references. PAO1 experiments were repeated with the PA14 medium conditions, and the same basic trends discussed in this paper still hold.

### Crystal violet biofilm assays

Biofilm assays were performed as previously described with minor modifications (50, 51). Briefly, culture inocula were grown in 100 µL of LB medium at 37 °C in a 96-well microtiter plate for ~16 h. Cultures were normalized and diluted ~1:100 in M63 medium with 1 mM magnesium sulfate and 0.4% arginine (instead of glucose and CAA). To a 96-well microtiter plate, 100 µL of the diluted culture was added to each well. Microtiter plates were then incubated at 37 °C for 24 h in a humidified environment to prevent culture evaporation. To remove unattached bacteria and spent medium, the microtiter dishes were inverted, then washed twice by gently immersing the plate in tap water followed by removing the liquid by briskly inverting the plate. Microtiter dish biofilms were stained by addition of 125 µL of 0.1% (w/v) crystal violet to each well, and incubation for 15 min at room temperature. After the crystal violet solution was removed, the plates were washed three times, as described above, with tap water. Plates were allowed to airdry overnight. The amount of crystal violet retained by each biofilm was measured by adding 150 µL of 30% (v/v) glacial acetic acid, incubating for 15 min at room temperature, and mixing by pipetting. Transfer of 100 µL of this mixture to a 96-well clear flat-bottom plate enabled spectrophotometric analysis at 550 nm. Each assay included 4 measurements (technical replicates), which were averaged, and the experiment was performed 5 times (biological replicates). The strains used in these assays are shown in Table S1. *P. aeruginosa* strains PAO1 and PA14 were initially described in (52) and (47), respectively. All clinical and environmental *P. aeruginosa* isolates were from the International *Pseudomonas* Consortium Database (IPCD) (43). These strains have both been phenotypically and genotypically characterized (44, 45).

### Flow cell experiments and data acquisition

Flow cells were prepared and inoculated as previously described (8) with the following modifications. Flow cells were purchased from two sources: Department of Systems Biology, Technical University of Denmark, and Ibidi (sticky-Slide *VI*^0.4^ with a glass coverslip). An in-line injection port (Ibidi) was used at the inlet for inoculating bacteria into the flow cell. For Ibidi flow cells, elbow connectors (Ibidi) were used to connect the chamber with tubing. The diluted bacteria culture was injected into the flow cell and allowed to incubate for 10-20 min without flow on the heating stage at 30 °C. Flow was then started at 3 mL/h for the entire acquisition time.

Images were taken using either an Andor iXon EMCCD camera with Andor IQ software on an Olympus IX81 microscope equipped with a Zero Drift Correction autofocus system or an Andor Neo sCMOS camera with Andor IQ software on an Olympus IX83 microscope equipped with a Zero Drift Correction 2 continuous autofocus system. Bright-field images were taken every 3 s (30 ms exposure time) on the IX81 system and every 100 ms (30 ms exposure time) on the IX83 system. For flagellum staining experiments, bright-field images were taken every 3 s (30 ms exposure time) on the IX81 system, and two fluorescence images (+0 and +1 μm above the imaging focal plane) were taken every 15 min (100 ms exposure time) using a Lambda LS (Sutter Instrument) xenon arc lamp and a GFP filter. On the IX81 system, total acquisition time was ~40 h, resulting in ~48000 images. On the IX83 system, total acquisition time was ~20 h, resulting in 720000 images. Image size was 67 μm × 67 μm (1024 × 1024 pixels).

### Multigenerational family tracking analysis

Image analysis, family tracking and manual validation, family tree plotting, and tree asymmetry calculations were performed in MATLAB as previously described (8) without modification. Fluorescence images were processed as follows to reduce noise and background signals and enhance flagella signals. Bandpass filtering, gamma correction, intensity percentile normalization, and then a green colormap were applied to the images. Fluorescence images were then overlaid on top of bright-field images using the lighten opacity setting. Probability distributions were obtained from the family trees as follows. The experimental probability distribution *P*_*n*_(*t*) is a 2D matrix, where the columns represent *n*, the number of cells present in one family, and each row is a time step *t*, the experimental image data acquisition interval (either every 3 s or 100 ms, depending on the data). For each time step *t* (in terms of lineage time, with each family starting at *t*_lineage_ = 0), we keep track of how many families have *n* = 0 cells, *n* = 1 cell, *n* = 2 cells, and so on. The proportion of families with *n* = 0, 1, 2, … cells then become one row in the matrix. This is equivalent to generating a histogram for *X*_*t*_(*ω*) using the states Σ as the bins. The full matrix is generated by repeating this for all time steps in the experimental data. Experimental moments were calculated by the formula 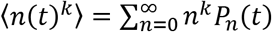, where *P*_*n*_(*t*) is the experimental probability distribution. MATLAB functions from the base installation of MATLAB R2015a, Statistics and Machine Learning Toolbox, Curve Fitting Toolbox, Image Processing Toolbox, Signal Processing Toolbox, and custom MATLAB functions were used for all analyses. In particular, the MATLAB functions “fit”, “fmincon”, and “ode45” were used for function fitting, nonlinear least-squares minimization with constraints, and numerical integration.

### Divide-detach stochastic model equations

Explanation of the model is given in the main text. The solution for the master equation (eq. 1), which is the model probability distribution *P*_*n*_(*t*), is given by

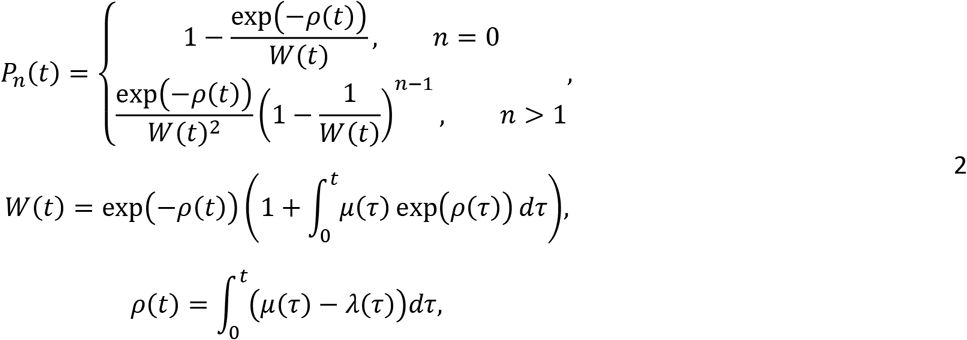

and *λ*(*t*) and *μ*(*t*) are the single cell division and detachment rates, respectively. Both rates are functions of time and positive (i.e., *λ*(*t*) > 0 and *μ*(*t*) > 0).

The first and second model moments are given by the following equations and the linear form of the rates, *λ*(*t*) = *L*_0_ + *L*_1_*t* and *μ*(*t*) = *C*_0_ + *C*_1_*t*, which are used for fitting the experimental moments

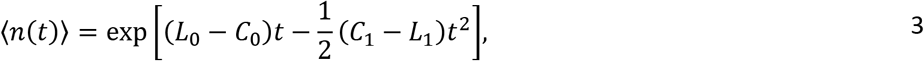

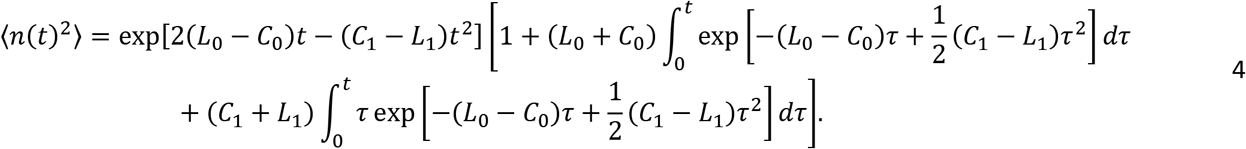

Evaluating the integral analytically in eq. 4 depends on the relative signs of {*L*_0_, *L*_1_, *C*_0_, *C*_1_}.

### Analytical solutions to the equations in the stochastic model

The solution for the master equation (eq. 1), which is the model probability distribution *P*_*n*_(*t*), can be found by using the so-called generating function

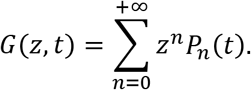

By plugging in the generating function into eq. 1, we obtain

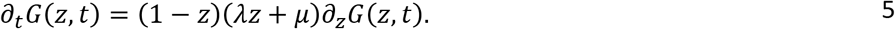

We can rewrite the previous equation in a Ricatti’s form, which reads

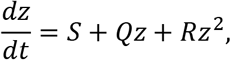

where *S* = −*μ*, *Q* = (*λ* + *μ*), and *R* = −*λ*. A particular solution of the previous equation is given by *Y*. Then the previous equation can be solved by quadrature *z*(*t*) = *x*(*t*) + *Y*(*t*) and

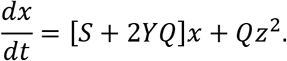

A change of variables 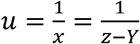 yields

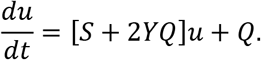

The solution of the Ricatti’s equation (53) is a homographic function

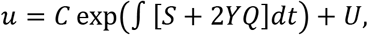

where *C* is an arbitrary constant and *U* is a particular solution. We can rewrite *u* as

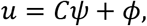

and the solution for *z* reads

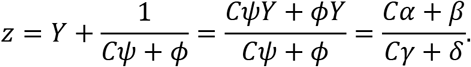

By using the Palm’s formulae (31–33, 35, 54), it is possible to find *P*_0_(*t*) and *P*_*n*_(*t*) as a function of *η*_*t*_ and *ξ*_*t*_, two unknown functions, which reads

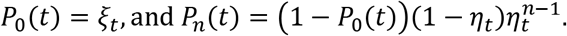

By means of geometric series, the generating function *G*(*z*, *t*) reads

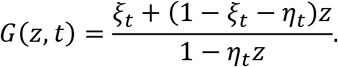

By plugging back this equation into eq. 5, we can find *ξ*_*t*_ and *η*_*t*_ and finally the solution (eq. 2).

To compare the model with the experimental results, we use the model moments defined as

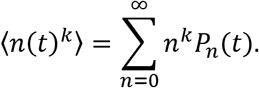

From the master equation (eq. 1), we can find

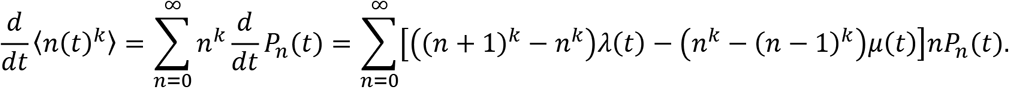

The first moment reads

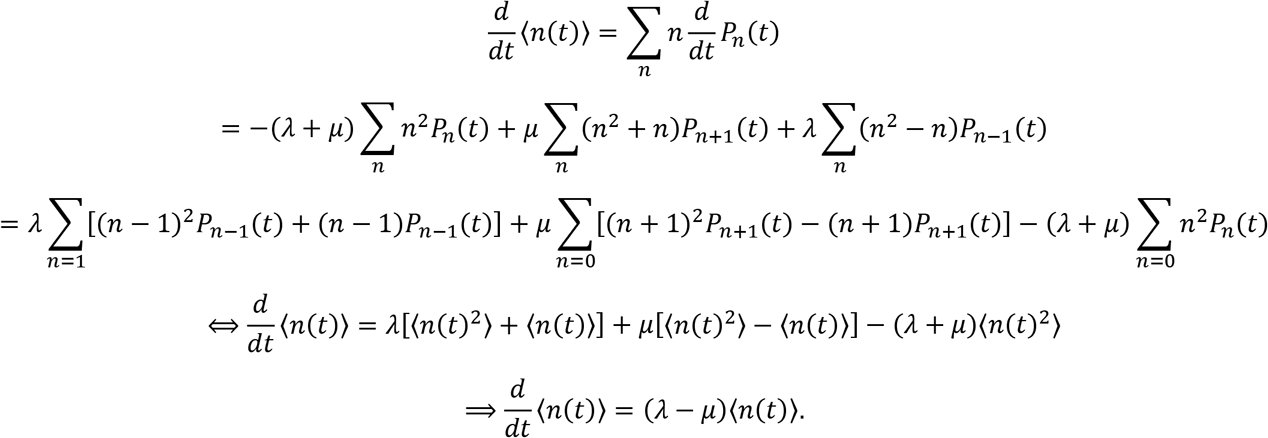

The solution to this differential equation is

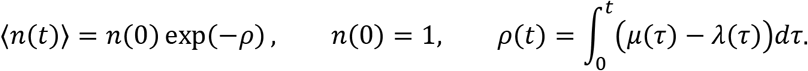

Plugging in the linear form of the rates, *λ*(*t*) = *L*_0_ + *L*_1_*t* and *μ*(*t*) = *C*_0_ + *C*_1_*t*, yields eq. 3.

The second moment (again, using the linear form of the rates, *λ*(*t*) = *L*_0_ + *L*_1_*t* and *μ*(*t*) = *C*_0_ + *C*_1_*t*) reads

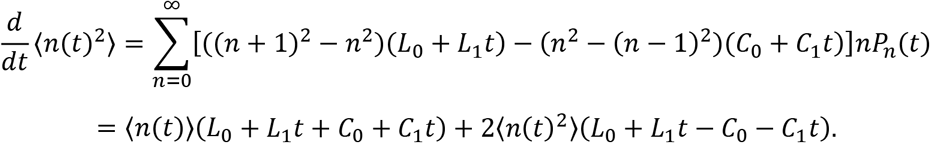

Elementary computations yield eq. 4.

### Hydrodynamic model of TFP retraction during flagellum-mediated spinning

We adapt the hydrodynamic model that we developed previously (37) to investigate the effects of TFP on flagellum-mediated spinning. Here, we consider a bacterium consisting of a cylindrical body attached to the surface at the pole opposite the flagellum, a helical filament for the flagellum of equal length to the body, and a straight filament 2/3 of the body length for a pilus protruding from the body. We use resistive force theory (55) to relate the angular velocities of each component of the bacterium to the torques from the flagellar motor, the viscous resistance from the fluid, and the flagellar hook which resists bending between the head and the flagellum. The resultant model is used to consider how TFP affect the angle that the bacterium makes with the surface during flagellum-mediated spinning and thereby “stand up” to a near vertical orientation commonly observed before detachment.

We use the example where the ratio of hook stiffness to motor torque is 0.5 to show the effects of TFP (see (37) for details of the stiffness/motor torque ratio). We show time using units of seconds and a torque value of 2 pN μm. We observe two significant effects on the surface angle when TFP retract during spinning: (i) the bacterium decreases its surface angle after retraction, (ii) the amplitude of oscillations in surface angle decreases after retraction. The opposite occurs when TFP extend during spinning: (i) the bacterium increases its surface angle after extension, (ii) the amplitude of oscillations in surface angle increases after extension. The strength of these effects depends on the choice of value of the flagellar motor torque and hook stiffness in the hydrodynamic model.

### TFP and flagellum activity metrics

To characterize appendage activity during family tree tracking, we use the algorithms previously described (8) for TFP activity and adapt them for flagellum activity. As previously described, TFP activity is inferred by recognizing surface translational motion, which is the predominant behavior for TFP-driven motion for *P. aeruginosa* cells that attach to the surface during early biofilm development. Analogous to this, the most common mode of flagellum activity is surface-attached “spinning,” where cells attach via one pole on the surface, and spin at angular velocities consistent with typical flagellum motor output (~5 rad/s) (36, 37). So, flagellum activity is inferred by recognizing surface rotational motion. Based on the majority of flagellum-mediated surface spinning behavior producing trajectories that are tightly clustered together and have strongly subdiffusive MSDs, the multi-parameter metric for flagellum activity is defined as follows. A bacterium has flagellum activity during a given time point when it is “spinning” and has non-zero displacement over a *w* frame moving window every *w*/10 frames. A cell that is “spinning” is defined as having the following characteristics during the *w* frame window: a Mean Squared Displacement (MSD) slope of less than 0.9 and having the maximum 2 point distance of its trajectory being greater than or equal to 15% of its maximum cell body length and less than its cell body length. As previously described, a value of *w* = 100 was used (8).

## Acknowledgements

CKL, GCLW, and MRP are supported by the Army Research Office (W911NF-18-1-0254). CKL, GCLW, and GAO are supported by the National Institutes of Health (1R01AI143730-01). KZ is supported by the National Key R&D Program of China (2018YFA0902102). RLT is supported by the Amgen Scholars U.S. Program.

We thank Joshua A. Keefe and Joshua S. Helali for their contributions to the initial family tree data validation during the early stages of this work.

## Supplementary Materials

**Figure S1.**
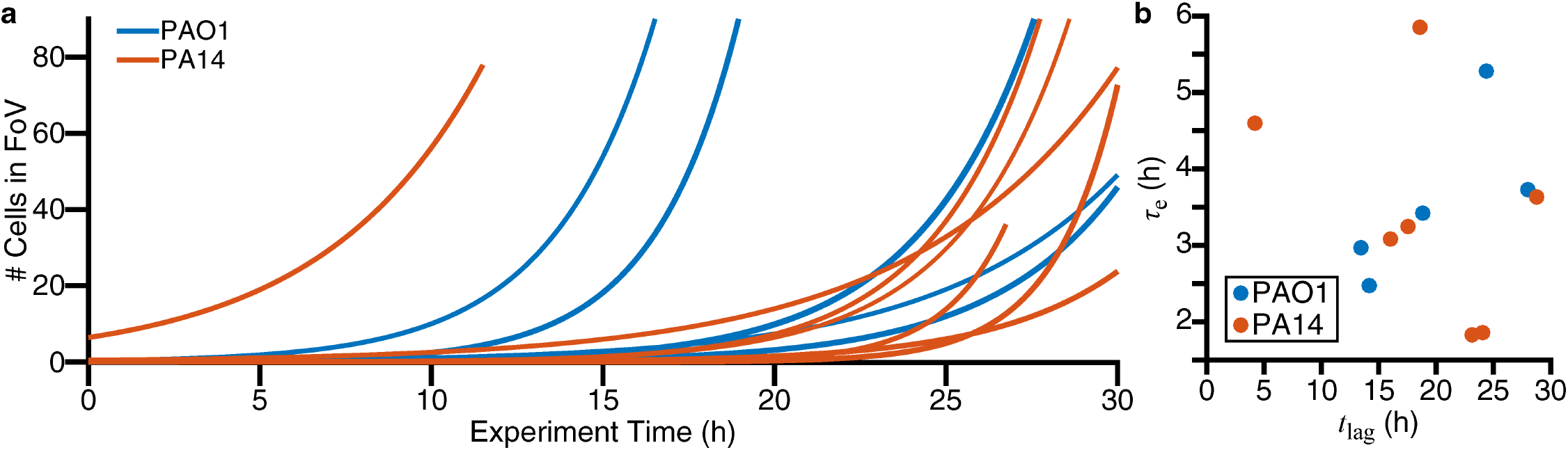
Exponential fits of bacteria count as a function of experiment time. (a) Each line is a fit to the function *N*(*t*) = *N*(0) exp))*t* − *t*_lag_)/*τ*_*e*_) for data shown in Figure 1. *N*(0) is the number of cells at time 0, *t*_lag_ characterizes the time scale of the lag period where *N* is roughly constant, and *τ*_*e*_ characterizes the time scale of exponential increase (1/*τ*_*e*_ characterizes the rate of exponential increase). (b) Plots of *τ*_*e*_ vs. *t*_lag_ for the fits in part (a).

**Figure S2.**
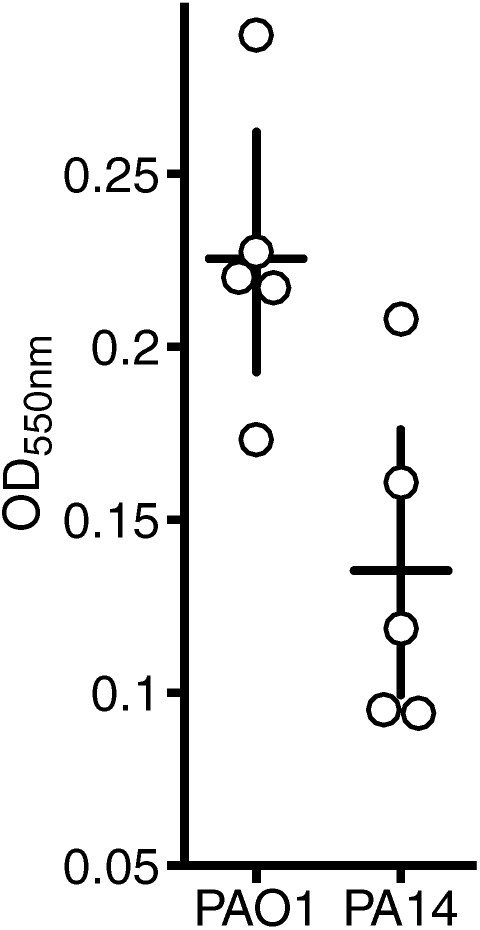
Monitoring biofilm formation via crystal violet assays. The OD_550nm_ values are proportional to the amount of biofilm stained by crystal violet. Circles represent individual biological replicates, each of which is the mean of 4 technical replicates. Horizontal lines represent the mean OD_550nm_ values. Vertical lines indicate the 95% confidence interval calculated from the bootstrap sampling distribution of the mean OD_550nm_ values.

**Figure S3.**
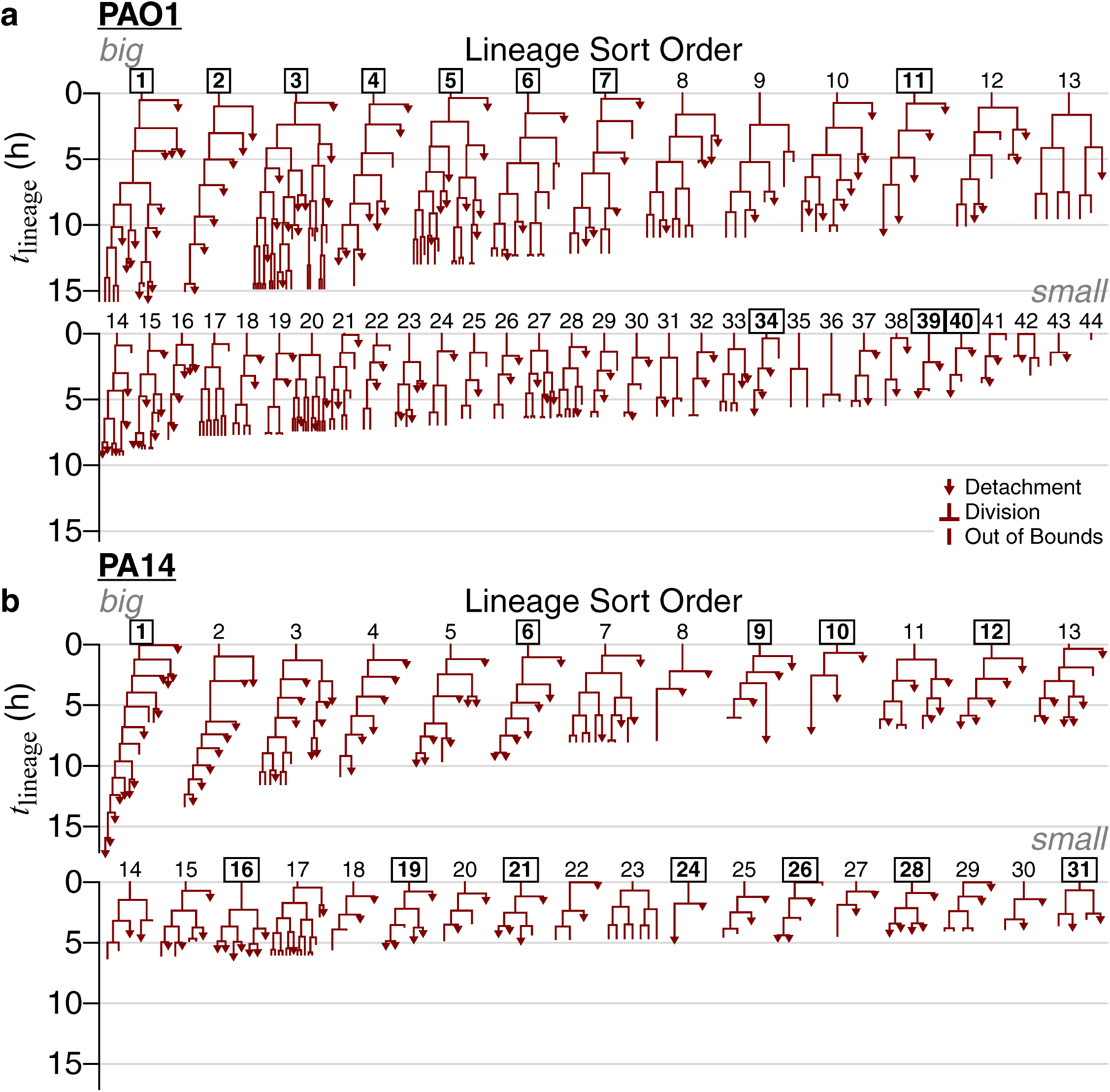
All families in the processive regime of reversible attachment, which is when cells divide at least once before detaching. We monitor the time that a given isolated family, consisting of an attached cell (founder cell) and its progeny (daughter cells), stays continually on the surface, which we designate as lineage time)*t* = *t*_lineage_). For each family, we begin tracking at the frame individual, founder bacteria attach and assign this as *t*_lineage_ = 0 h. We continue tracking until either the entire family detaches, or until we lose track of that family (where we can no longer distinguish individual cells, or the cells move out of the recording boundaries). This final time point is recorded as the family’s residence time. For families that we lose track of, their true residence times can be greater than or equal to these recorded values, while for families that end in detachment, their recorded and true residence times are equal. Only families that end in detachment are used for calculating the average residence time to avoid the uncertainty in actual residence times for families that we lose track of. For each regime and strain, we sort families by residence times in descending order, which sorts them by the amount of time that they have continuously contacted the surface. Lineage indices that are boxed and bolded are the families selected for use in the model. The criteria used to select families for the model are described as follows. We set the minimum number of families required for the model at ≈10 families. First, we select all families that result in lineage “extinction” events (where we observe the family ending in a detachment event), because these families all have defined outcomes (recorded and true residence times are equal). Conversely, families that we lose track of (where we can no longer distinguish individual cells, or the cells move out of the recording boundaries) have undefined outcomes (true residence times are greater than or equal to the recorded residence times). For PA14, this results in 12 selected families (#1, 6, 9, 10, 12, 16, 19, 21, 24, 26, 28, and 31), which meets the minimum family number requirement. For PAO1, this results in 4 selected families (#11, 34,39, and 40), which is not enough. To increase the number of selected families for PAO1, we apply a residence time cutoff on the families with undefined outcomes, so that all selected families have defined outcomes at least in the lineage time window the analysis is performed on. A residence time cutoff of *t*_lineage_ = 12 h results in an additional 7 families (#1-7) for a total of 11 selected families for PAO1, which meets the minimum family number requirement.

**Figure S4.**
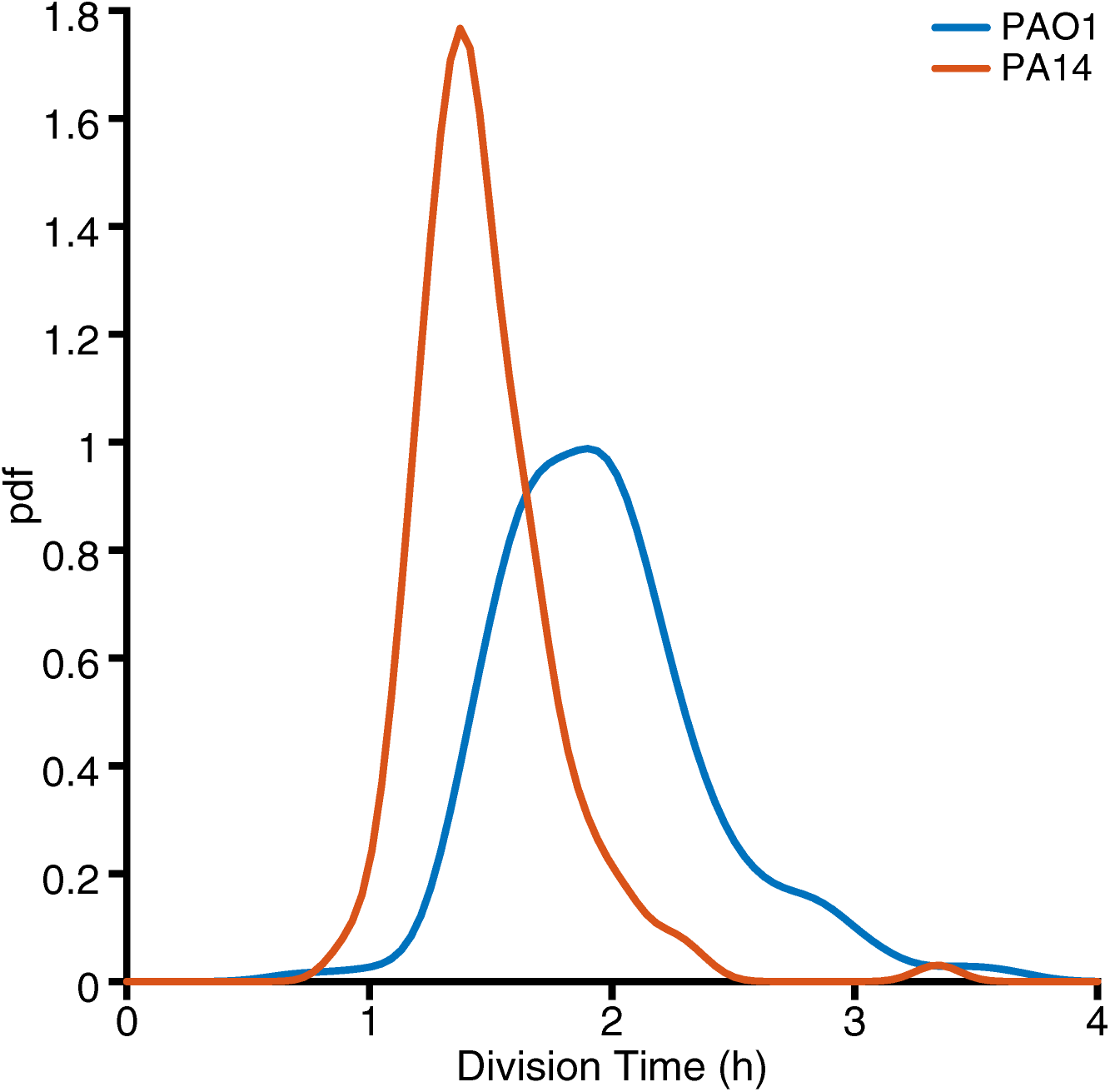
Distribution of division times for PAO1 and PA14. Division time is calculated as the time between consecutively observed division events in a family. Distributions are plotted as pdf (probability density function) calculated via kernel density estimation. PAO1 has a median division time of 1.92 h with a 95% confidence interval of (1.87 h, 1.96 h), and PA14 has a median division time of 1.42 h with a 95% confidence interval of (1.38 h, 1.45 h). PAO1 has a mean ± standard deviation division time of 1.96 ± 0.44 h, and PA14 has a mean ± standard deviation division time of 1.48 ± 0.30 h.

**Figure S5.**
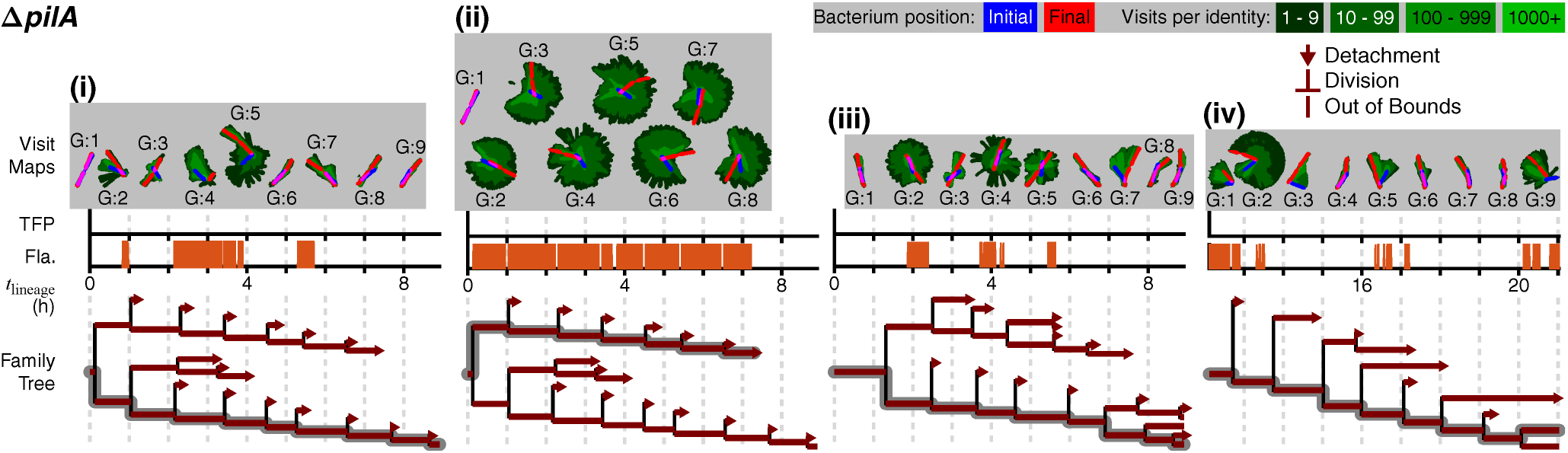
Appendage activity tracking for the Δ*pilA* mutant. We repeat the analysis in Figure 8 for the Δ*pilA* mutant and find one predominant behavior up the 9 generations shown here, consistent with this strain having predominantly one-legged division branching (8). We observe no TFP activity, which is consistent with this strain having no TFP; we observe sporadic, but prolonged, flagellum activity, which is consistent with observations of this strain spinning on the surface for prolonged periods of time; and we observe detachment events without flagellum activity, which is consistent with observations of detachment events where cells did not have a labeled flagellum. Beyond generation 9, the Δ*pilA* mutant continues to have mainly one-legged division-branching for multiple subsequent generations.

**Figure S6.**
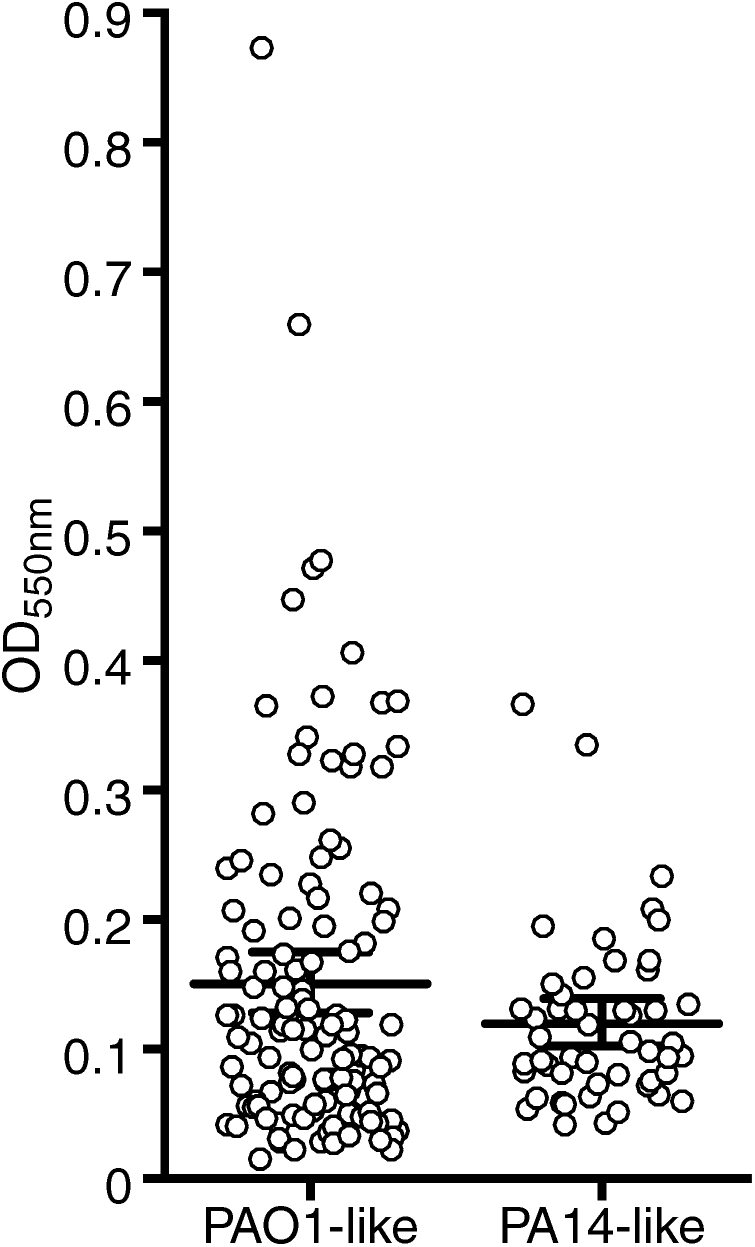
Crystal violet biofilm assay results for 35 *P. aeruginosa* strains (25 PAO1-like and 10 PA14-like strains, including PAO1 and PA14 strains) in the International *P. aeruginosa* Consortium Database (IPCD). These strains are identified as either PAO1-like or PA14-like based on their phylogeny (i.e., same phylogenetic sub-group as either PAO1 or PA14) (42–45). The OD_550nm_ values are proportional to the amount of biofilm stained by crystal violet. Circles represent individual biological replicates, each of which is the mean of 4 technical replicates. Longer horizontal lines represent the mean OD_550nm_ values. Vertical lines and error bars indicate the 95% confidence interval calculated from the bootstrap sampling distribution of the mean OD_550nm_ values. Comparing these distributions shows that the mean OD_550nm_ value for the PAO1-like strains are higher than the mean OD_550nm_ value for PA14-like strains (p-value of 0.02).

**Table S1.**
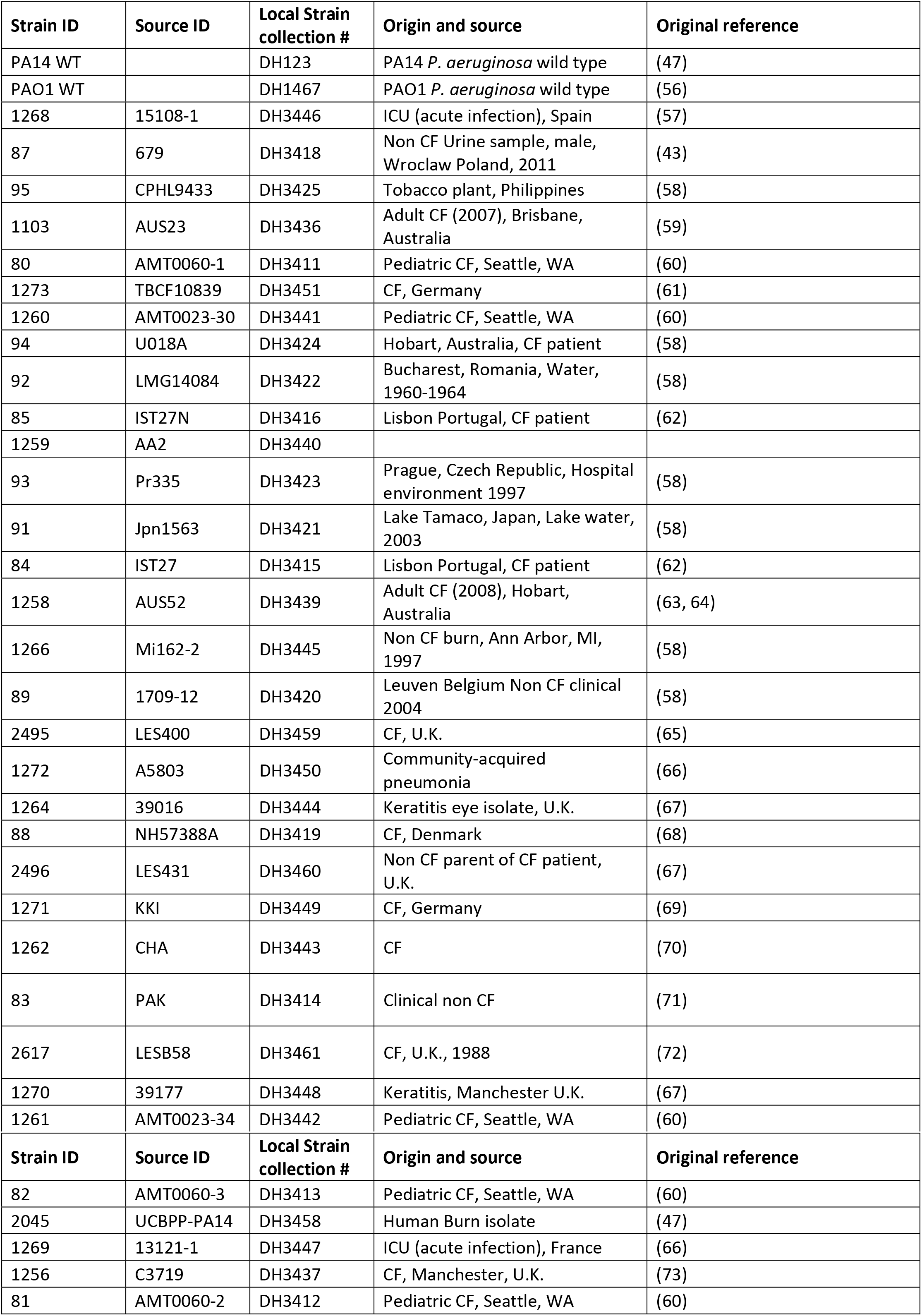
Strains used in the crystal violet biofilm assays. The collection of isolates was described in (43). (CF = cystic fibrosis)

## References

1. Singh PK, Schaefer AL, Parsek MR, Moninger TO, Welsh MJ, Greenberg EP. 2000. Quorum-sensing signals indicate that cystic fibrosis lungs are infected with bacterial biofilms. Nature 407:762–764.

2. Hall-Stoodley L, Costerton JW, Stoodley P. 2004. Bacterial biofilms: from the Natural environment to infectious diseases. Nature Reviews Microbiology 2:95.

3. Biteen JS, Blainey PC, Cardon ZG, Chun M, Church GM, Dorrestein PC, Fraser SE, Gilbert JA, Jansson JK, Knight R, Miller JF, Ozcan A, Prather KA, Quake SR, Ruby EG, Silver PA, Taha S, van den Engh G, Weiss PS, Wong GCL, Wright AT, Young TD. 2016. Tools for the Microbiome: Nano and Beyond. ACS Nano 10:6–37.

4. O’Toole GA, Wong GCL. 2016. Sensational Biofilms: Surface Sensing in Bacteria. Curr Opin Microbiol 30:139–146.

5. Hug I, Deshpande S, Sprecher KS, Pfohl T, Jenal U. 2017. Second messenger–mediated tactile response by a bacterial rotary motor. Science 358:531–534.

6. McCarter L, Hilmen M, Silverman M. 1988. Flagellar Dynamometer Controls Swarmer Cell Differentiation of *V. parahaemolyticus*. Cell 54:345–351.

7. Ellison CK, Dalia TN, Vidal Ceballos A, Wang JC-Y, Biais N, Brun YV, Dalia AB. 2018. Retraction of DNA-bound type IV competence pili initiates DNA uptake during natural transformation in *Vibrio cholerae*. Nature Microbiology 3:773–780.

8. Lee CK, de Anda J, Baker AE, Bennett RR, Luo Y, Lee EY, Keefe JA, Helali JS, Ma J, Zhao K, Golestanian R, O’Toole GA, Wong GCL. 2018. Multigenerational memory and adaptive adhesion in early bacterial biofilm communities. Proceedings of the National Academy of Sciences 115:4471–4476.

9. Guvener ZT, Harwood CS. 2007. Subcellular Location Characteristics of the *Pseudomonas aeruginosa* GGDEF Protein, WspR, Indicate that It Produces Cyclic-di-GMP in Response to Growth on Surfaces. Mol Microbiol 66:1459–1473.

10. O’Connor JR, Kuwada NJ, Huangyutitham V, Wiggins PA, Harwood CS. 2012. Surface Sensing and Lateral Subcellular Localization of WspA, the Receptor in a Chemosensory-like System Leading to C-di-GMP Production. Mol Microbiol 86:720–729.

11. Chen Y, Chai Y, Guo JH, Losick R. 2012. Evidence for cyclic Di-GMP-mediated signaling in *Bacillus subtilis*. Journal of bacteriology 194:5080–5090.

12. Ferreira RB, Chodur DM, Antunes LC, Trimble MJ, McCarter LL. 2012. Output Targets and Transcriptional Regulation by a Cyclic Dimeric GMP-Responsive Circuit in the *Vibrio parahaemolyticus* Scr Network. J Bacteriol 194:914–924.

13. Gomelsky M, Galperin MY. 2013. Bacterial second messengers, cGMP and c-di-GMP, in a quest for regulatory dominance. The EMBO journal 32:2421–2423.

14. Romling U, Galperin MY, Gomelsky M. 2013. Cyclic di-GMP: the first 25 years of a universal bacterial second messenger. Microbiology and molecular biology reviews: MMBR 77:1–52.

15. Jones CJ, Utada A, Davis KR, Thongsomboon W, Zamorano Sanchez D, Banakar V, Cegelski L, Wong GCL, Yildiz FH. 2015. C-di-GMP Regulates Motile to Sessile Transition by Modulating MshA Pili Biogenesis and Near-Surface Motility Behavior in Vibrio cholerae. PLOS Pathogens 11:e1005068.

16. Zamorano-Sánchez D, Xian W, Lee CK, Salinas M, Thongsomboon W, Cegelski L, Wong GCL, Yildiz FH. 2019. Functional Specialization in *Vibrio cholerae* Diguanylate Cyclases: Distinct Modes of Motility Suppression and c-di-GMP Production. mBio 10:e00670–19.

17. Luo Y, Zhao K, Baker AE, Kuchma SL, Coggan KA, Wolfgang MC, Wong GCL, O’Toole GA. 2015. A Hierarchical Cascade of Second Messengers Regulates *Pseudomonas aeruginosa* Surface Behaviors. mBio 6:1–11.

18. Persat A, Inclan YF, Engel JN, Stone HA, Gitai Z. 2015. Type IV Pili Mechanochemically Regulate Virulence Factors in *Pseudomonas aeruginosa*. Proc Natl Acad Sci U S A 112:7563–7568.

19. Costerton JW, Stewart PS, Greenberg EP. 1999. Bacterial biofilms: a common cause of persistent infections. Science 284:1318–22.

20. Huangyutitham V, Güvener ZT, Harwood CS. 2013. Subcellular Clustering of the Phosphorylated WspR Response Regulator Protein Stimulates Its Diguanylate Cyclase Activity. mBio 4:e00242–13.

21. Armbruster CR, Lee CK, Parker-Gilham J, de Anda J, Xia A, Zhao K, Tseng BS, Hoffman LR, Jin F, Harwood CS, Wong GCL, Parsek MR. 2019. Heterogeneity in surface sensing suggests a division of labor in *Pseudomonas aeruginosa* populations. eLife 8:e45084.

22. Irie Y, Borlee BR, O’Connor JR, Hill PJ, Harwood CS, Wozniak DJ, Parsek MR. 2012. Self-produced exopolysaccharide is a signal that stimulates biofilm formation in *Pseudomonas aeruginosa*. Proc Natl Acad Sci U S A 109:20632–20636.

23. Zhao K, Tseng BS, Beckerman B, Jin F, Gibiansky ML, Harrison JJ, Luijten E, Parsek MR, Wong GCL. 2013. Psl Trails Guide Exploration and Microcolony Formation in *Pseudomonas aeruginosa* Biofilms. Nature (London, U K) 497:388–391.

24. Colvin KM, Gordon VD, Murakami K, Borlee BR, Wozniak DJ, Wong GC, Parsek MR. 2011. The pel polysaccharide can serve a structural and protective role in the biofilm matrix of *Pseudomonas aeruginosa*. PLoS Pathog 7:e1001264.

25. Sauer K, Camper AK, Ehrlich GD, Costerton JW, Davies DG. 2002. *Pseudomonas aeruginosa* Displays Multiple Phenotypes during Development as a Biofilm. Journal of Bacteriology 184:1140–1154.

26. Davey ME, Caiazza NC, O’Toole GA. 2003. Rhamnolipid Surfactant Production Affects Biofilm Architecture in *Pseudomonas aeruginosa* PAO1. Journal of Bacteriology 185:1027–1036.

27. Henrici AT. 1933. Studies of Freshwater Bacteria: I. A Direct Microscopic Technique. J Bacteriol 25:277–87.

28. Zobell CE, Allen EC. 1935. The Significance of Marine Bacteria in the Fouling of Submerged Surfaces. Journal of Bacteriology 29:239–251.

29. Sanchez A, Golding I. 2013. Genetic Determinants and Cellular Constraints in Noisy Gene Expression. Science 342:1188–1193.

30. Shiryayev AN. 2012. Selected Works of AN Kolmogorov: Volume II Probability Theory and Mathematical Statistics, vol 26. Springer Science & Business Media.

31. Arley N. 1949. On the “birth-and-death” process. Scandinavian Actuarial Journal 1949:21–26.

32. Kendall DG. 1949. Stochastic Processes and Population Growth. Journal of the Royal Statistical Society Series B (Methodological) 11:230–282.

33. Kendall DG. 1948. On the Generalized “Birth-and-Death” Process. Ann Math Statist 19:1–15.

34. Novozhilov AS, Karev GP, Koonin EV. 2006. Biological applications of the theory of birth-and-death processes. Briefings in Bioinformatics 7:70–85.

35. Feller W. On the Theory of Stochastic Processes, with Particular Reference to Applications, p 403–432. In (ed), University of California Press,

36. Conrad JC, Gibiansky ML, Jin F, Gordon VD, Motto Da, Mathewson Ma, Stopka WG, Zelasko DC, Shrout JD, Wong GCL. 2011. Flagella and Pili-Mediated Near-Surface Single-Cell Motility Mechanisms in *P. aeruginosa*. Biophys J 100:1608–1616.

37. Bennett RR, Lee CK, De Anda J, Nealson KH, Yildiz FH, O’Toole GA, Wong GCL, Golestanian R. 2016. Species-Dependent Hydrodynamics of Flagellum-Tethered Bacteria in Early Biofilm Development. J RSoc, Interface 13:20150966.

38. Gibiansky ML, Conrad JC, Jin F, Gordon VD, Motto DA, Mathewson MA, Stopka WG, Zelasko DC, Shrout JD, Wong GCL. 2010. Bacteria Use Type IV Pili to Walk Upright and Detach from Surfaces. Science (Washington, DC, U S) 330:197.

39. Coates J, Park BR, Le D, Şimşek E, Chaudhry W, Kim M. 2018. Antibiotic-induced population fluctuations and stochastic clearance of bacteria. eLife 7:e32976.

40. Irie Y, Roberts AEL, Kragh KN, Gordon VD, Hutchison J, Allen RJ, Melaugh G, Bjarnsholt T, West SA, Diggle SP. 2017. The *Pseudomonas aeruginosa* PSL Polysaccharide Is a Social but Noncheatable Trait in Biofilms. mBio 8:e00374–17.

41. Nadell CD, Drescher K, Foster KR. 2016. Spatial structure, cooperation and competition in biofilms. Nature Reviews Microbiology 14:589–600.

42. Freschi L, Jeukens J, Kukavica-Ibrulj I, Boyle B, Dupont M-J, Laroche J, Larose S, Maaroufi H, Fothergill JL, Moore M, Winsor GL, Aaron SD, Barbeau J, Bell SC, Burns JL, Camara M, Cantin A, Charette SJ, Dewar K, Déziel É, Grimwood K, Hancock REW, Harrison JJ, Heeb S, Jelsbak L, Jia B, Kenna DT, Kidd TJ, Klockgether J, Lam JS, Lamont IL, Lewenza S, Loman N, Malouin F, Manos J, McArthur AG, McKeown J, Milot J, Naghra H, Nguyen D, Pereira SK, Perron GG, Pirnay J-P, Rainey PB, Rousseau S, Santos PM, Stephenson A, Taylor V, Turton JF, Waglechner N, et al. 2015. Clinical utilization of genomics data produced by the international *Pseudomonas aeruginosa* consortium. Frontiers in Microbiology 6:1036.

43. De Soyza A, Hall AJ, Mahenthiralingam E, Drevinek P, Kaca W, Drulis-Kawa Z, Stoitsova SR, Toth V, Coenye T, Zlosnik JEA, Burns JL, Sá-Correia I, De Vos D, Pirnay J-P, J. Kidd T, Reid D, Manos J, Klockgether J, Wiehlmann L, Tümmler B, McClean S, Winstanley C, pathogens” EFfCABCsvdocf. 2013. Developing an international *Pseudomonas aeruginosa* reference panel. MicrobiologyOpen 2:1010–1023.

44. Cullen L, Weiser R, Olszak T, Maldonado RF, Moreira AS, Slachmuylders L, Brackman G, Paunova-Krasteva TS, Zarnowiec P, Czerwonka G, Reilly J, Drevinek P, Kaca W, Melter O, De Soyza A, Perry A, Winstanley C, Stoitsova SR, Lavigne R, Mahenthiralingam E, Sá-Correia I, Coenye T, Drulis-Kawa Z, Augustyniak D, Valvano MA, McClean S. 2015. Phenotypic characterization of an international *Pseudomonas aeruginosa* reference panel: strains of cystic fibrosis (CF) origin show less in vivo virulence than non-CF strains. Microbiology 161:1961–1977.

45. Freschi L, Bertelli C, Jeukens J, Moore MP, Kukavica-Ibrulj I, Emond-Rheault J-G, Hamel J, Fothergill JL, Tucker NP, McClean S, Klockgether J, de Soyza A, Brinkman FSL, Levesque RC, Winstanley C. 2018. Genomic characterisation of an international *Pseudomonas aeruginosa* reference panel indicates that the two major groups draw upon distinct mobile gene pools. FEMS Microbiology Letters 365.

46. Mischel W, Ebbesen EB, Raskoff Zeiss A. 1972. Cognitive and attentional mechanisms in delay of gratification. Journal of Personality and Social Psychology 21:204–218.

47. Rahme LG, Stevens EJ, Wolfort SF, Shao J, Tompkins RG, Ausubel FM. 1995. Common virulence factors for bacterial pathogenicity in plants and animals. Science 268:1899–1902.

48. Kuchma SL, Ballok AE, Merritt JH, Hammond JH, Lu W, Rabinowitz JD, O’Toole GA. 2010. Cyclic-di-GMP-mediated repression of swarming motility by *Pseudomonas aeruginosa*: the *pilY1* gene and its impact on surface-associated behaviors. J Bacteriol 192:2950–64.

49. de Anda J, Lee EY, Lee CK, Bennett RR, Ji X, Soltani S, Harrison MC, Baker AE, Luo Y, Chou T, O’Toole GA, Armani AM, Golestanian R, Wong GCL. 2017. High-Speed “4D” Computational Microscopy of Bacterial Surface Motility. ACS Nano 11:9340–9351.

50. O’Toole GA, Kolter R. 1998. Initiation of biofilm formation in *Pseudomonas fluorescens* WCS365 proceeds via multiple, convergent signalling pathways: a genetic analysis. Molecular Microbiology 28:449–461.

51. O’Toole GA. 2011. Microtiter dish biofilm formation assay. J Vis Exp doi:10.3791/2437:e2437.

52. Holloway BW. 1955. Genetic Recombination in *Pseudomonas aeruginosa*. Microbiology 13:572–581.

53. Soare MV, Teodorescu PP, Toma I. 2007. Ordinary differential equations with applications to mechanics, vol 585. Springer.

54. Khinchin AY, Andrews D, Quenouille MH. 2013. Mathematical methods in the theory of queuing. Courier Corporation.

55. Gray J, Hancock GJ. 1955. The Propulsion of Sea-Urchin Spermatozoa. J Exp Biol 32:802–814.

## Supplementary References

56. Stover CK, Pham XQ, Erwin AL, Mizoguchi SD, Warrener P, Hickey MJ, Brinkman FS, Hufnagle WO, Kowalik DJ, Lagrou M, Garber RL, Goltry L, Tolentino E, Westbrock-Wadman S, Yuan Y, Brody LL, Coulter SN, Folger KR, Kas A, Larbig K, Lim R, Smith K, Spencer D, Wong GK, Wu Z, Paulsen IT, Reizer J, Saier MH, Hancock RE, Lory S, Olson MV. 2000. Complete genome sequence of *Pseudomonas aeruginosa* PAO1, an opportunistic pathogen. Nature 406:959–964.

57. Köhler T, Buckling A, van Delden C. 2009. Cooperation and virulence of clinical *Pseudomonas aeruginosa* populations. Proceedings of the National Academy of Sciences 106:6339–6344.

58. Pirnay J-P, Bilocq F, Pot B, Cornelis P, Zizi M, Van Eldere J, Deschaght P, Vaneechoutte M, Jennes S, Pitt T, De Vos D. 2009. *Pseudomonas aeruginosa* Population Structure Revisited. PLOS ONE 4:e7740.

59. O’Carroll MR, Syrmis MW, Wainwright CE, Greer RM, Mitchell P, Coulter C, Sloots TP, Nissen MD, Bell SC. 2004. Clonal strains of *Pseudomonas aeruginosa* in paediatric and adult cystic fibrosis units. European Respiratory Journal 24:101–106.

60. Mulcahy LR, Burns JL, Lory S, Lewis K. 2010. Emergence of *Pseudomonas aeruginosa* Strains Producing High Levels of Persister Cells in Patients with Cystic Fibrosis. Journal of Bacteriology 192:6191–6199.

61. Bezuidt OK, Klockgether J, Elsen S, Attree I, Davenport CF, Tümmler B. 2013. Intraclonal genome diversity of *Pseudomonas aeruginosa* clones CHA and TB. BMC Genomics 14:416.

62. Leitão JH, Alvim T, Sá-Correia I. 1996. Ribotyping of *Pseudomonas aeruginosa* isolates from patients and water springs and genome fingerprinting of variants concerning mucoidy. Pathogens and Disease 13:287–292.

63. Bradbury R, Champion A, Reid DW. 2008. Poor clinical outcomes associated with a multi-drug resistant clonal strain of *Pseudomonas aeruginosa* in the Tasmanian cystic fibrosis population. Respirology 13:886–892.

64. Kidd TJ, Ramsay KA, Hu H, Marks GB, Wainwright CE, Bye PT, Elkins MR, Robinson PJ, Rose BR, Wilson JW, Grimwood K, Bell SC. 2013. Shared *Pseudomonas aeruginosa* genotypes are common in Australian cystic fibrosis centres. European Respiratory Journal 41:1091–1100.

65. Salunkhe P, Smart CHM, Morgan JAW, Panagea S, Walshaw MJ, Hart CA, Geffers R, Tümmler B, Winstanley C. 2005. A Cystic Fibrosis Epidemic Strain of *Pseudomonas aeruginosa* Displays Enhanced Virulence and Antimicrobial Resistance. Journal of Bacteriology 187:4908–4920.

66. Wiehlmann L, Wagner G, Cramer N, Siebert B, Gudowius P, Morales G, Köhler T, van Delden C, Weinel C, Slickers P, Tümmler B. 2007. Population structure of *Pseudomonas aeruginosa*. Proceedings of the National Academy of Sciences 104:8101–8106.

67. Stewart RMK, Wiehlmann L, Ashelford KE, Preston SJ, Frimmersdorf E, Campbell BJ, Neal TJ, Hall N, Tuft S, Kaye SB, Winstanley C. 2011. Genetic Characterization Indicates that a Specific Subpopulation of *Pseudomonas aeruginosa* Is Associated with Keratitis Infections. Journal of Clinical Microbiology 49:993–1003.

68. Hoffmann N, Rasmussen TB, Jensen P, Stub C, Hentzer M, Molin S, Ciofu O, Givskov M, Johansen HK, Høiby N. 2005. Novel Mouse Model of Chronic *Pseudomonas aeruginosa* Lung Infection Mimicking Cystic Fibrosis. Infection and Immunity 73:2504–2514.

69. Cramer N, Wiehlmann L, Ciofu O, Tamm S, Høiby N, Tümmler B. 2012. Molecular Epidemiology of Chronic *Pseudomonas aeruginosa* Airway Infections in Cystic Fibrosis. PLOS ONE 7:e50731.

70. Toussaint B, Delicattree I, Vignais PM. 1993. *Pseudomonas aeruginosa* Contains an IHF-like Protein That Binds to the *algD* Promoter. Biochemical and Biophysical Research Communications 196:416–421.

71. Totten PA, Lory S. 1990. Characterization of the type a flagellin gene from *Pseudomonas aeruginosa* PAK. Journal of Bacteriology 172:7188–7199.

72. Kukavica-Ibrulj I, Bragonzi A, Paroni M, Winstanley C, Sanschagrin F, O’Toole GA, Levesque RC. 2008. In Vivo Growth of *Pseudomonas aeruginosa* Strains PAO1 and PA14 and the Hypervirulent Strain LESB58 in a Rat Model of Chronic Lung Infection. Journal of Bacteriology 190:2804–2813.

73. Jones AM, Dodd ME, Doherty CJ, Govan JR, Webb AK. 2002. Increased treatment requirements of patients with cystic fibrosis who harbour a highly transmissible strain of *Pseudomonas aeruginosa*. Thorax 57:924–925.

